# What a running template needs: leg damping and a hip torque

**DOI:** 10.64898/2026.11.13.766384

**Authors:** M. Mert Ankarali, Uluç Saranlı

## Abstract

A template, the simplest model that reproduces a behaviour, is a hypothesis about what the body must do. For running, the spring-loaded inverted pendulum (SLIP) captures the bounce of the centre of mass but not the force behind it: its ground reaction force (GRF) is symmetric in time and always points along the leg. Its many extensions have never faced a common test. We fitted 34 templates to stance GRF and ranked them by the Bayesian information criterion and cross-validation on six open human datasets (414 runner–speed conditions, 2–5 m s^−1^, treadmill and overground) and on aerial running in two guinea-fowl datasets. Every dataset agreed. A running template needs two elements: leg damping, without which errors stayed above 0.3 body weights (BW), and a force off the leg axis, without which the fore-aft error stayed near that of SLIP. In humans, SLIP with leg damping and a hip torque that decays through stance (TD-SLIP) has both; its two-parameter form was best in the three largest datasets and within 9.3 BIC per condition of the best elsewhere (errors 0.15–0.18 BW). Guinea fowl needed the same two elements in another form, a virtual pivot above the centre of mass, and kept it on slippery ground. Parameters varied between runners far more than their gaits, and overground running softened the leg by 8% without changing the ranking: runners share the template’s structure, not its numbers. The GRF fixes the template but not how leg and hip share the work, which joint-level data must decide.

**Summary statement:** Across 34 models, six human datasets and running guinea fowl, a running template needs leg damping and a hip torque; individuals share its structure, not its numbers.

## Introduction

Full and Koditschek (1999) proposed templates and anchors as neuromechanical hypotheses of legged locomotion (Fig. 1). A template is the simplest model, with the fewest variables and parameters, that exhibits a targeted behaviour; an anchor is a more realistic model, grounded in morphology and physiology (legs, joint torques, muscles and neural circuits), that explains how the body manages its many degrees of freedom and energy so that it behaves like the template (Full and Koditschek, 1999; Holmes et al., 2006). Animals with two, four, six or eight legs and very different postures bounce like a spring–mass system (Blickhan and Full, 1993). For running, the spring-loaded inverted pendulum (SLIP), a point mass bouncing on a massless spring leg, is the canonical template of running in both biomechanics and robotics (Geyer and Saranlı, 2018): it captures the centre-of-mass (COM) trajectory, the stance time and the self-stability of running across species (Blickhan, 1989; Geyer et al., 2006; McMahon and Cheng, 1990). Its ground reaction force (GRF), however, is too symmetric: SLIP is conservative, so its stance trajectory is symmetric in time about mid-stance (Saranlı et al., 2010), its force always points along the leg, and it cannot produce the early vertical peak, the skewed force profile or the fore-aft impulses of real running (Maus et al., 2015; Srinivasan and Holmes, 2008). How far runners depart from the spring–mass ideal is itself a runner trait: elite runners are closer to a symmetric bounce than recreational runners, with a shorter lag between the fore-aft braking–propulsion transition and the vertical loading–unloading transition and a smaller error of the best-fitting spring–mass vertical force (Burns et al., 2023).

**Figure 1:**
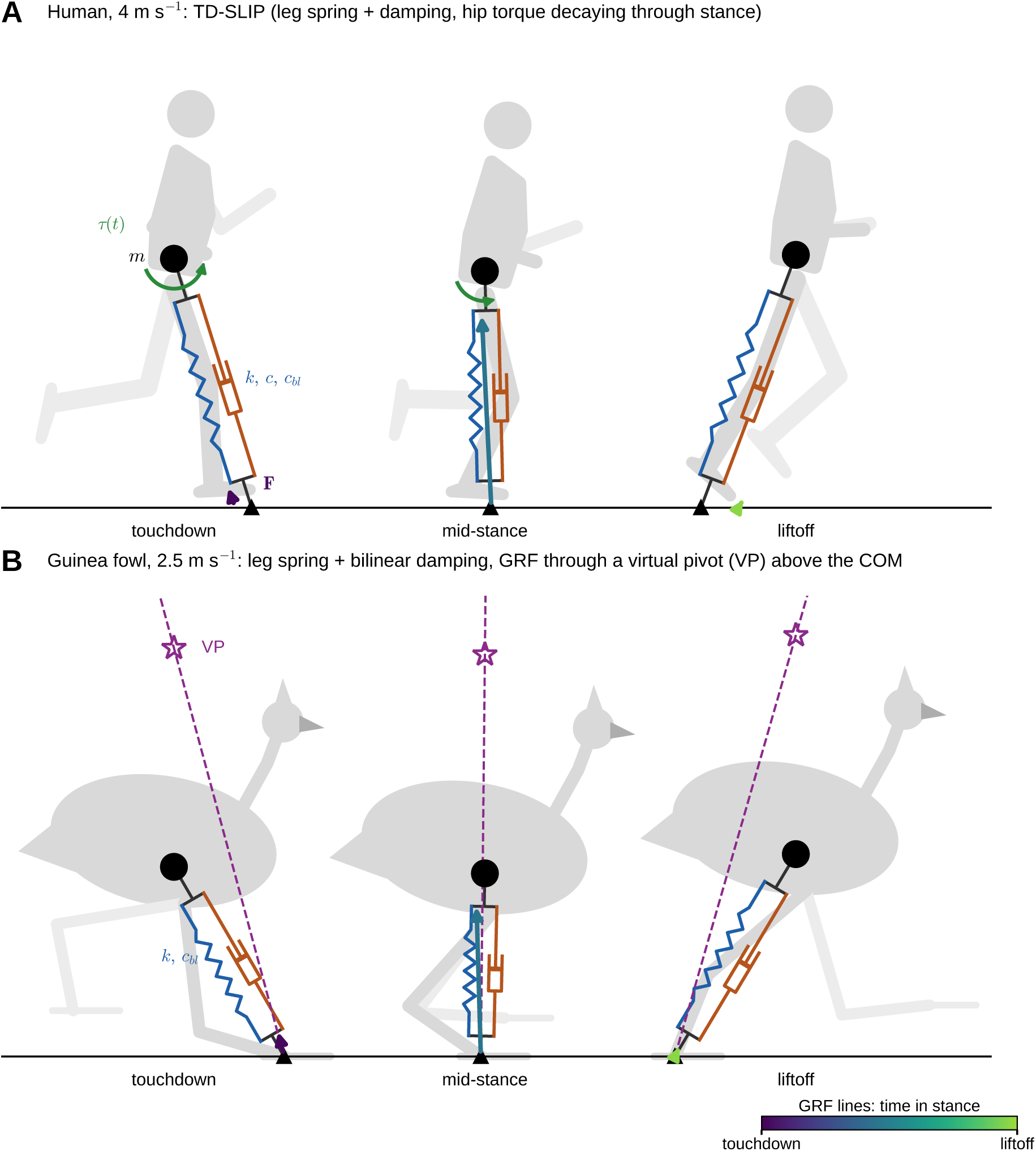
Templates on running animals. (A) A human runner (participant 1 of Van Hooren and Meijer (2024), 4 m s^−1^) at touchdown, mid-stance and liftoff of one stance, with the TD-SLIP family of templates drawn on top: a point mass at the COM (black disc) on a massless spring–damper leg (spring *k*, linear and bilinear damping *c*, *c_bl_*) to a fixed foot (triangle), and a hip torque *τ* (*t*) that decays through stance (green arc). Body from the motion-capture markers (sagittal projection; arms and head drawn schematically). (B) A guinea fowl (Daley and Birn-Jeffery, 2026b) in one steady aerial-running step (2.5 m s^−1^), with the template preferred by the bird data: a spring leg with bilinear damping whose force passes through a virtual pivot (VP, star) above the COM; COM and feet from the data, body schematic. Arrows, measured GRF at the centre of pressure (stance foot in the bird), coloured from touchdown to liftoff. Lengths are scaled by leg length *l*_0_.

Templates are not only descriptions of animals; they have shaped how legged machines are built and controlled. Raibert’s running machines separated control of a springy leg into hopping height, forward speed and body attitude (Raibert, 1986). The hexapod RHex, built with compliant legs and driven by an open-loop clock, runs with the COM dynamics of SLIP (Altendorfer et al., 2001; Saranlı et al., 2001). The biped ATRIAS — “Assume The Robot Is A Sphere” — was designed with light legs and series springs so that its own dynamics approximate a spring–mass system, and its controllers exploit the interplay between this tuned passive compliance and actuation to walk and run robustly (Hubicki et al., 2016, 2018). In template-based control the template becomes a literal control target: a controller makes a machine with many degrees of freedom behave like the template, gait-level commands such as speed and hop height are given to the template, and the remaining degrees of freedom, above all body pitch, are stabilised by the embedding controller. This turned the open-loop, pitch-unstable pronking of an underactuated hexapod into a stable gait with explicit speed and height control (Ankaralı and Saranlı, 2011); SLIP can be imposed as the hybrid zero dynamics of an asymmetric hopper (Poulakakis and Grizzle, 2009); and hierarchical controllers embed pendular COM templates at the virtual COM of floating-torso robots while a separate loop holds body orientation, adapting to the control authority of each contact mode (De et al., 2022). With a hip torque driven by a clock, one template family spans walking, trotting and running trots of a quadruped and the transitions between them, on hardware (Akbaba et al., 2025). Gait control on the template relies on analytic approximations of its stance map (Geyer and Saranlı, 2018; Saranlı et al., 2010), which predicted the position and velocity of a physical monopod with average errors of about 2% and 7% once its parameters had been identified (Uyanık et al., 2015). Control rules found on biomechanical templates, such as swing-leg retraction (Seyfarth et al., 2003) and virtual-pendulum posture control (Sharbafi et al., 2013), were likewise formulated as laws that a machine can implement. The template is thus the interface through which biology informs robotics, and the question of which template is right belongs to both fields: a template that matches animal forces with the right mechanical elements tells a designer which elements a machine needs to run like an animal, and robotics in turn shows what each element can do when it is controlled rather than passive.

Many additions have been proposed. Leg damping and actuation along the leg (linear, non-linear and stiffness-switching laws) break this symmetry and reshape the vertical force (Abraham et al., 2015; Ludwig et al., 2012; Saranlı et al., 2010). In legged robots with an actuator in series with the leg spring, damping is not only a loss: the actuator can make the leg behave like a lossy SLIP whose damping coefficient is a control input, so that energy is removed or added along the leg throughout stance while the analytic approximations of the template stay valid (Seçer and Saranlı, 2014, 2018). Hip torques — prescribed in time (Ankaralı and Saranlı, 2010; Shen et al., 2014), driven by a clock (Seipel and Holmes, 2007), or produced by a hip spring — make the force non-radial; virtual pivot points (VPP) above or below the COM redirect the force towards a fixed point on the trunk (Drama et al., 2020; Maus et al., 2010), and a VPP improves the GRF predicted by full-body musculoskeletal models of walking (Wagner et al., 2024); the central pivot point (CPP) routes it through the COM (Karagöz et al., 2020; Sever et al., 2026); and foot or leg masses reproduce the impact transient (Clark et al., 2014). Each was introduced with its own data and fitting method. Which of these elements is *necessary*, which is merely sufficient, and whether the answer depends on the runners or the laboratory, have not been tested on a common footing.

The choice matters beyond describing forces. Templates serve as the plants of task-level models of sensorimotor control: a unicycle template of the running cockroach showed that antennal wall following needs feedback of the rate of approach as well as of distance (Cowan et al., 2006), and that the antenna, reconfigured passively by locomotion and contact mechanics, simplifies this control (Mongeau et al., 2013). In such closed-loop analyses behaviour emerges from sensing, neural processing and body mechanics together (Cowan et al., 2014), and neural signals cannot be interpreted without the mechanics they drive (Tytell et al., 2011). A stance template plays the same role for stride-to-stride control of running: a template that assigns the force to the wrong mechanical element will assign it to the wrong controller as well.

A runner is not a low-order, linear or time-invariant system, and its forces vary from stance to stance, between speeds, between individuals and between laboratories. A template is useful only if it is parsimonious and robust to this biological variability: it should describe the force with few parameters and keep describing it when the runner, the speed or the laboratory changes. Choosing such a model is a model-selection problem — goodness of fit traded against complexity, and tested on data the model has not seen (Ankaralı et al., 2014). A feature that is real is not necessarily worth modelling: human walking dynamics are measurably left–right asymmetric, yet a model that ignores the asymmetry predicts held-out strides as well or better and has much smaller parameter uncertainty — walking is symmetric enough (Ankaralı et al., 2015). Variability between individuals raises a second question. Glass knifefish differ about twofold in the stiffness and damping of their locomotor plant, yet track a moving refuge within about 15–20% of one another, because feedback compensates for the mismatch (Uyanik et al., 2020). A template may therefore be shared by all runners while its parameters differ between them; the test of a template is whether its structure transfers to new runners. We built the benchmark around this logic: the information criterion penalizes parameters, cross-validation holds out speeds and runners, and replication across datasets and protocols holds out the laboratory.

TD-SLIP, a SLIP with leg damping and a hip torque that decays linearly in time, was proposed in earlier work, which showed on digitized GRF curves that it fits running much better than SLIP (Ankaralı et al., 2012); it has since served as the abstraction for the design and control of small running robots (Truax et al., 2024). Here we ask the general question: which template explains the GRF of human running, does the answer replicate across datasets, and does it hold for a running bird? We implemented 34 templates in five families (Fig. 2), fitted each to every runner–speed condition of six open human datasets and to aerial running in two guinea-fowl datasets, one of them on high- and low-friction ground, under the same periodic-gait constraints, and ranked them with the Bayesian information criterion (BIC) and cross-validation. We also asked what fixes which force component, how the fitted parameters change between treadmill and overground running, and what the measured force-line geometry implies for a template’s hip torque. We use the effective-footprint picture (Srinivasan and Holmes, 2008) throughout, because it shows directly what a hip torque does to the force (Fig. 3).

**Figure 2:**
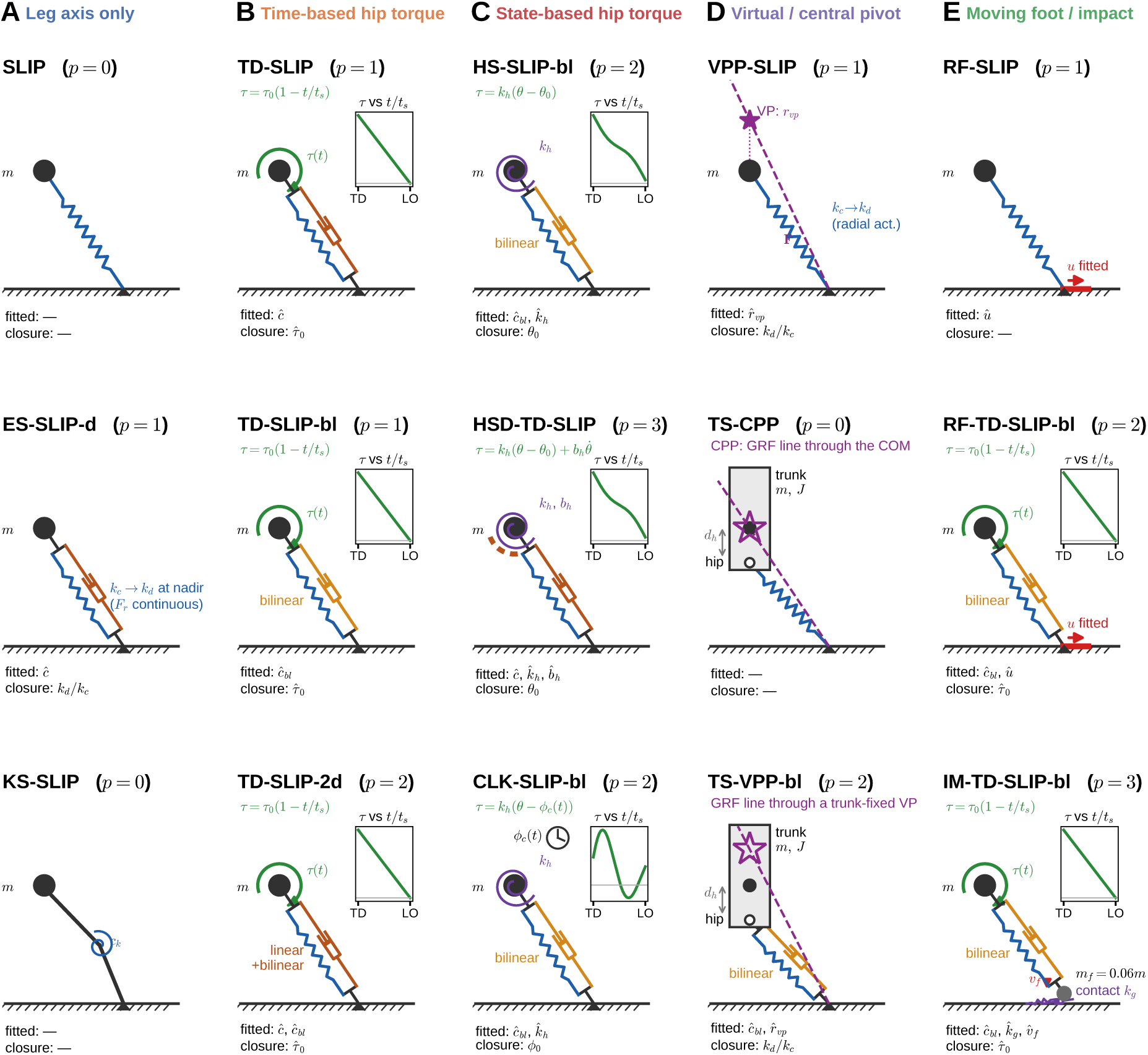
The template ladder: three representative members of each family. (A) Leg axis only: SLIP, linear damping with a force-continuous stiffness switch, two-segment knee leg. (B) Time-based hip torque: TD-SLIP with linear, bilinear and linear+bilinear damping. (C) State-based hip torque: hip spring, hip spring–damper on TD-SLIP, clock-torqued SLIP. (D) Pivots: point-mass VPP, trunk with the central pivot point (CPP), trunk with a fixed VPP. (E) Moving foot and impact: foot moving at a fitted speed, with and without the ramp torque, and a foot mass with a soft contact. *p*, number of shape parameters fitted to the GRF; insets show the torque profile (TD, touchdown; LO, liftoff). All 34 templates are shown in Figs S1–S3.

**Figure 3:**
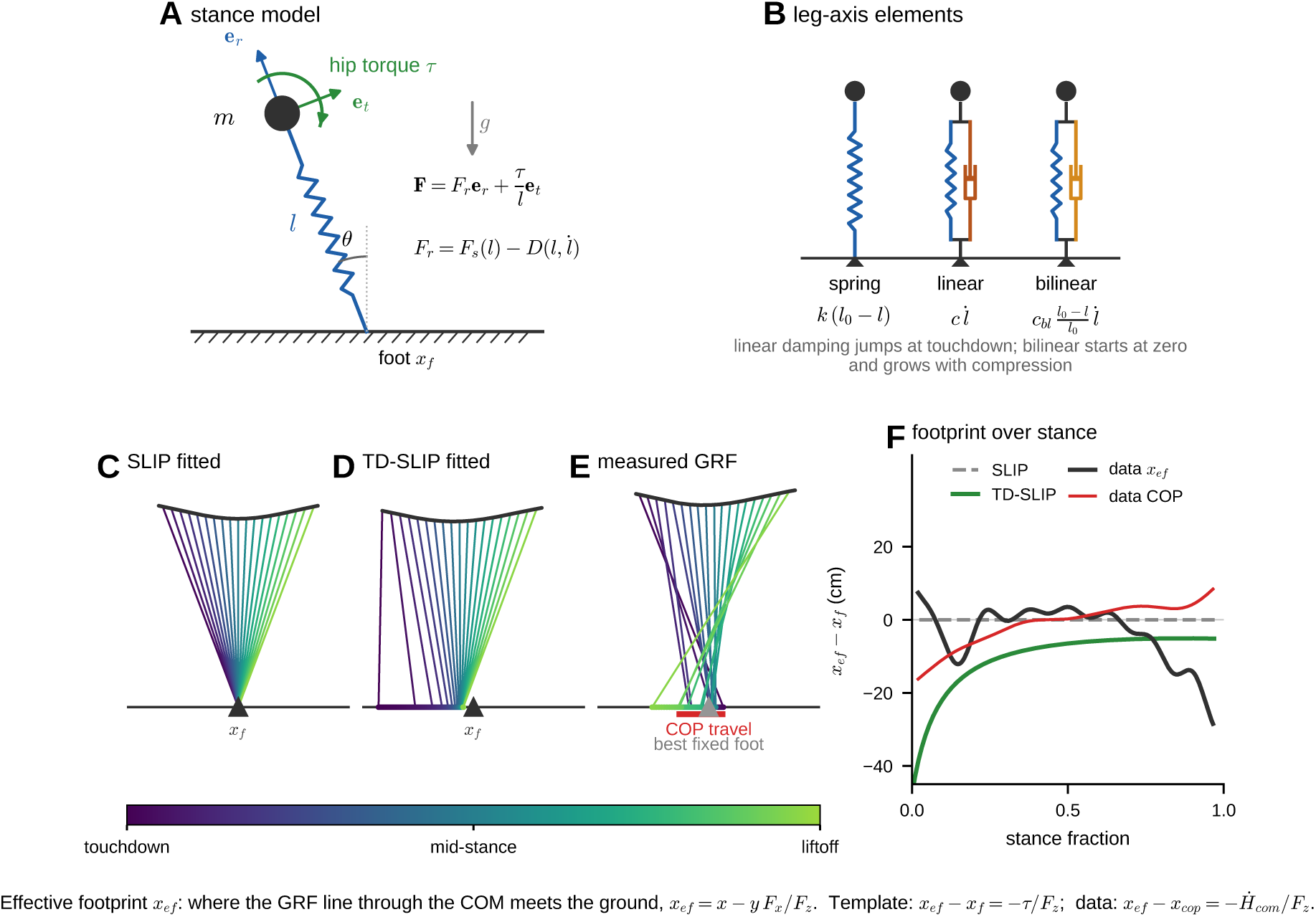
Stance model and the effective footprint. (A) The GRF of every template is a radial force *F_r_* along the leg plus the hip torque *τ* divided by leg length across it. (B) Leg-axis elements: spring, linear damper (force jump at touchdown) and bilinear damper (zero at touchdown, growing with compression). (C–F) Effective footprints for runner 1 of Van Hooren and Meijer (2024) at 4 m s^−1^. In C–E the GRF line through the COM (black path) is drawn to the ground at 19 instants, coloured from touchdown to liftoff; dots on the ground are the effective footprints *x_ef_* (*F_z_ >* 0.2 BW). (C) SLIP fitted to this condition: the force is radial and every line passes through the foot (triangle). (D) TD-SLIP fitted to the same condition: the hip torque shifts each line by −*τ/F_z_*, far behind the foot early in stance when *F_z_* is small. (E) Measured GRF, relative to the best fixed foot; the red bar marks COP travel. (F) The same footprints over stance, relative to the foot (templates) or to the best fixed foot (data): SLIP stays at zero, the TD-SLIP footprint approaches the foot from behind, and the measured footprint stays near a fixed point through mid-stance while the COP rolls forward beneath it.

## Materials and methods

### Data

Six open datasets of level human running were used (Table S1): Van Hooren and Meijer (2024) (19 runners, split-belt treadmill, 2.78–5 m s^−1^), Fukuchi et al. (2017) (39 runners, treadmill, 2.5–4.5 m s^−1^), Carter et al. (2024) (50 runners, treadmill, at habitual speed and at 3.33 m s^−1^), Baker et al. (2025) (15 runners, treadmill and overground), and Riglet et al. (2024) (30 runners, overground). The digitized curves of Kram et al. (1998) and Hamill et al. (1983) used in the 2012 study were refitted for reproduction. Another candidate dataset (Wang et al., 2025) was excluded because its trials are trimmed to the plate contacts, so flight time and body mass cannot be recovered. Raw data were downloaded from the public repositories and are not redistributed.

For a bird, we used the guinea-fowl (*Numida meleagris*) ensemble of Daley and Birn-Jeffery (2026b) (13 birds, level overground running; 500 Hz; data deposited by Daley and Birn-Jeffery, 2026a). Its force plates give the whole-body net GRF, so only aerial-running steps, in which one leg is on the ground, can be compared with a one-leg template; we used the steady aerial-running steps of the authors’ analysis sample. Stance was the interval with vertical force above 0.1 body weights, and stances whose duration differed by more than 25% from the authors’ marker-based contact time (merged across a short flight) were rejected. The lower foot marker served as the COP, and the COM height at touchdown came from the authors’ reconstructed COM. Steps were grouped by bird and dimensionless speed 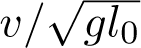 (bins at 1.3, 1.6 and 1.9), giving 37 conditions with at least five steps (435 stances, 1.4–4.0 m s^−1^). Because birds placed the fitted virtual pivot near the upper end of the grid used for humans, the pivot height was searched up to 2*l*_0_ for this dataset.

A second guinea-fowl dataset (Goldsmith et al., 2026) (7 birds, 1.6–2.2 kg; data deposited by Daley et al., 2026) recorded straight runs and turns over a force-plate runway on high-friction (control) and low-friction (slippery) ground; we used the straight runs (60 and 70 trials). The export gives the whole-body net GRF and the COP at 800 Hz, already filtered, together with the authors’ COM estimate. As above, only stances bounded by force below 0.1 body weights (flight of at least 8 ms) entered, and only steady steps: forward speed changing by less than 10% from touchdown to the next touchdown and mean vertical force over the step within 15% of body weight. This dataset has a measured COP, so it tests the pivot geometry without the foot-marker proxy. Its exported COM height at touchdown was about 1.6 hip heights, against 1.06 in the first dataset from the same laboratory with birds of the same size, which points to a constant offset; we shifted each bird’s COM height by one constant (common to both surfaces) to 1.06 hip heights and repeated the key fits with the exported height. Steps were grouped by bird, surface and dimensionless speed (bins at 1.1, 1.3 and 1.6), giving 13 conditions on each surface (225 and 247 stances, 1.3–2.5 m s^−1^).

Forces were low-pass filtered (4th-order zero-lag Butterworth, 50 Hz, or left unfiltered when sampled at 120 Hz). A stance phase was the contiguous interval with vertical force above 50 N; steps with implausible duration, a peak below 1.3–1.5 body weights (BW), contact on a second plate, or missing flight phase were rejected. The centre of pressure (COP) was expressed in the belt frame for treadmills. Each runner–speed condition was reduced to a condition-mean stance (time normalised to stance fraction, averaged and rescaled to the mean stance time); in total 414 human and 63 guinea-fowl conditions entered the analysis (Table S1). The COM trajectory during stance was reconstructed from the GRF under three steady-state assumptions (zero net fore-aft impulse, equal height and vertical velocity after the flight phase, and zero net angular impulse about the COM over stance), with the COM height at touchdown taken from pelvis markers or the body model where available (Table S1). The reconstructed horizontal COM offset agreed with the pelvis markers within 0.9 ± 1.7 cm on Van Hooren and Meijer (2024) and within 0.4 cm on a Fukuchi condition.

### Templates

All templates share a point mass *m* on a massless leg from a foot point to the COM (Fig. 3A):

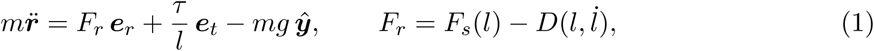

where *l* is leg length, ***e****_r_* and ***e****_t_* the radial and tangential unit vectors, *F_s_* the leg spring, *D* the leg damping and *τ* the hip torque. Liftoff occurs when *F_r_* returns to zero. The 34 templates combine one element from each of the following (Fig. 2; full list in Table S2 and Figs S1–S3):

- *leg spring*: linear *k*(*l*_0_−*l*), power law, a two-segment leg with a knee spring, or a stiffness switch at maximum compression (with or without a force jump; Ludwig et al. 2012);
- *leg damping*: none, linear *cl̇*, bilinear 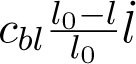 (Abraham et al., 2015), both, or compression-only;
- *hip law*: none; time-based torques (linearly decaying ramp *τ*_0_(1 −*t/t_s_*) of TD-SLIP, constant, half-stance, half-sine, force-proportional); state-based torques (a hip torsional spring *k_h_*(*θ −θ*_0_), a hip spring–damper, or a clock-torqued SLIP in which the spring tracks a phase *ϕ*(*t*); Seipel and Holmes 2007);
- *pivot*: a virtual pivot on the point mass with one or two offsets, or a rigid trunk whose hip torque routes the GRF through the CPP (the COM) or a trunk-fixed VPP (Karagöz et al., 2020; Maus et al., 2010);
- *foot*: fixed, moving at a fitted speed, moving with the measured COP, or a foot mass (6% of body mass) with a Hunt–Crossley contact (Hunt and Crossley, 1975).

The templates are grouped in five families for presentation: leg axis only, time-based hip torque, state-based hip torque, virtual or central pivot, and moving foot (Fig. 2). Model names combine the elements; for example TD-SLIP-2d is the ramp-torque template with linear plus bilinear damping, HSD-TD-SLIP the TD-SLIP with an added hip spring–damper, and TS-CPP the trunk model with the CPP.

### Gait constraints and parameters

For each condition the touchdown angle, touchdown velocity, dimensionless stiffness *k*^^^ = *kl*_0_*/mg* and one closure parameter were solved so that the simulated stance satisfies five periodic-gait constraints taken from the data, as in Ankaralı et al. (2012): equal fore-aft velocity at touchdown and liftoff; return to the touchdown height and vertical velocity after the measured flight time; the measured stance time; and the measured average speed. The closure parameter closes the energy balance of non-conservative templates (for example the torque amplitude *τ*^_0_ = *τ*_0_*/mgl*_0_ of TD-SLIP or the rest angle of a hip spring). The remaining dimensionless *shape* parameters (0–3 per template; for example the damping constants 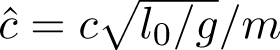 were fitted to the GRF; only these are counted as free parameters *p*. Equations were integrated with a fourth-order Runge–Kutta scheme (Δ*t* = 0.2 ms). Leg length *l*_0_ was the COM-to-COP distance at touchdown.

### Fitting, information criteria and cross-validation

The error measure was the mean squared error of fore-aft and vertical force in BW over stance. For every condition and template the error was tabulated on a grid of the shape parameters (1-D: 15–41 nodes; 2-D: 9–21 per axis; 3-D: 11 ×7 ×7), each node requiring a constrained gait solve. Fits minimised the error over grid nodes; on Van Hooren and Meijer (2024) the single-condition fits were further refined by a bounded Nelder–Mead search on the re-simulated error. For each condition, the BIC was computed with Gaussian errors of unknown variance and an effective sample size *n* = 2*f_c_t_s_* per force component (*f_c_* = 50 Hz; with *f_c_* = 20 Hz the top three were unchanged and the rank correlation over all templates was 0.98). We report the BIC excess per condition relative to the best template of each dataset, which is comparable across datasets of different size.

We chose the BIC over the Akaike information criterion (AIC) because the aim is the simplest adequate template, not the best predictor of the force. No template is the true model of a runner— all models are wrong (Box and Draper, 1987) — but the two criteria answer different questions. The BIC penalty grows with the number of observations, which makes it consistent: if a model of finite dimension generated the data, the BIC would select it with probability approaching one as data accumulate (Schwarz, 1978). The AIC estimates the expected information loss (Akaike, 1974), is asymptotically equivalent to leave-one-out cross-validation, and keeps adding parameters as data accumulate (Shao, 1997). Between simplicity and fit, the benchmark prefers simplicity. The AIC and its small-sample correction (AICc; Hurvich and Tsai, 1989) were computed for comparison.

Cross-validation complements the BIC rather than repeating it. It tests a template on data not used to fit it, needs no count of parameters — equal weighting of all parameters is itself an assumption, since a damping constant and a torque amplitude need not cost the same— and remains meaningful when the Gaussian error model behind the information criteria is wrong (Arlot and Celisse, 2010). Its outcome depends on how the data are split (Shao, 1993); we therefore did not split at random but held out the units that a template is expected to generalise to. Cross-validation tested shared shape parameters on held-out data: a held-out speed of the same runner, a held-out runner at the same speed, and one parameter set for all conditions tested on a held-out runner. Gait observables of the held-out condition (speed, stance and flight time) were always used, so the templates predict force *shape*. Medians of test errors are reported because a shared parameter set sometimes has no periodic solution for a held-out condition. For a held-out runner this occurred in at most 1% of tests for all templates except the damped trunk CPP template (7%) and the two-segment leg (21%); for a held-out speed it was more frequent for some multi-parameter templates (up to 48% for the undamped hip spring–damper and 19% for HSD-TD-SLIP).

### Additional analyses

*Component-only fits* fitted the fore-aft or the vertical force alone to find the best error each template can reach on each component. *Impact*: the foot-mass template was fitted to the 19 conditions at 4 m s^−1^. *Low-pass control*: the same 19 conditions were low-pass filtered at 15 Hz to remove the impact transient and refitted with eight key templates. *Treadmill versus overground*: in Baker et al. (2025) each runner’s overground condition was paired with the same runner’s treadmill condition at the nearest speed (median difference 0.00 m s^−1^) and fitted parameters compared with Wilcoxon signed-rank tests; across datasets, Riglet et al. (2024) (overground) was compared with the three treadmill datasets with COP data over the same dimensionless speed range 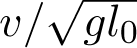, using a linear speed term and runner-clustered bootstrap intervals. *Force-line geometry*: effective footprints *x_ef_* = *x −y F_x_/F_z_* were computed from the reconstructed COM and measured GRF; the measured pivot point was the point minimising the distance to all force lines; and template inversion computed the hip torque and leg-axis force that a point-mass template on a fixed foot would need to reproduce the measured GRF exactly. *Foot strike*: fitted damping constants were compared between the foot-strike categories reported by Fukuchi et al. (2017).

### Software and use of generative AI

All analyses used Python (NumPy, SciPy, Numba, pandas, Matplotlib). Code development, data processing and drafting of the text were assisted by a generative AI system (Claude, Anthropic); all analyses, numbers and text were checked and approved by the authors, who take full responsibility for the content.

## Results

### The effective footprint separates the template families

For any point-mass template the line of the GRF through the COM meets the ground at the effective footprint, and *x_ef_ −x_f_* = −*τ/F_z_* (Fig. 3). A force along the leg passes through the foot, so every force line of SLIP and of all leg-axis templates converges on the foot (Fig. 3C); a hip torque moves the line off the foot in proportion to *τ/F_z_* (Fig. 3D,F). Measured force lines do neither. In a single runner the effective footprint stays within a few centimetres of one point through mid-stance while the COP rolls forward beneath it, and moves backward near touchdown and toe-off (Fig. 3E,F). Averaged over runners, it lies behind the touchdown COP for the first 15% of stance, sits 0.15–0.27 m ahead of it through mid-stance and swings back near toe-off (Fig. 4A). By the moment identity *x_ef_ −x_cop_* = −*Ḣ /F_z_* the mid-stance plateau follows the COP, while the early and late excursions reflect the rate of change of angular momentum about the COM.

**Figure 4:**
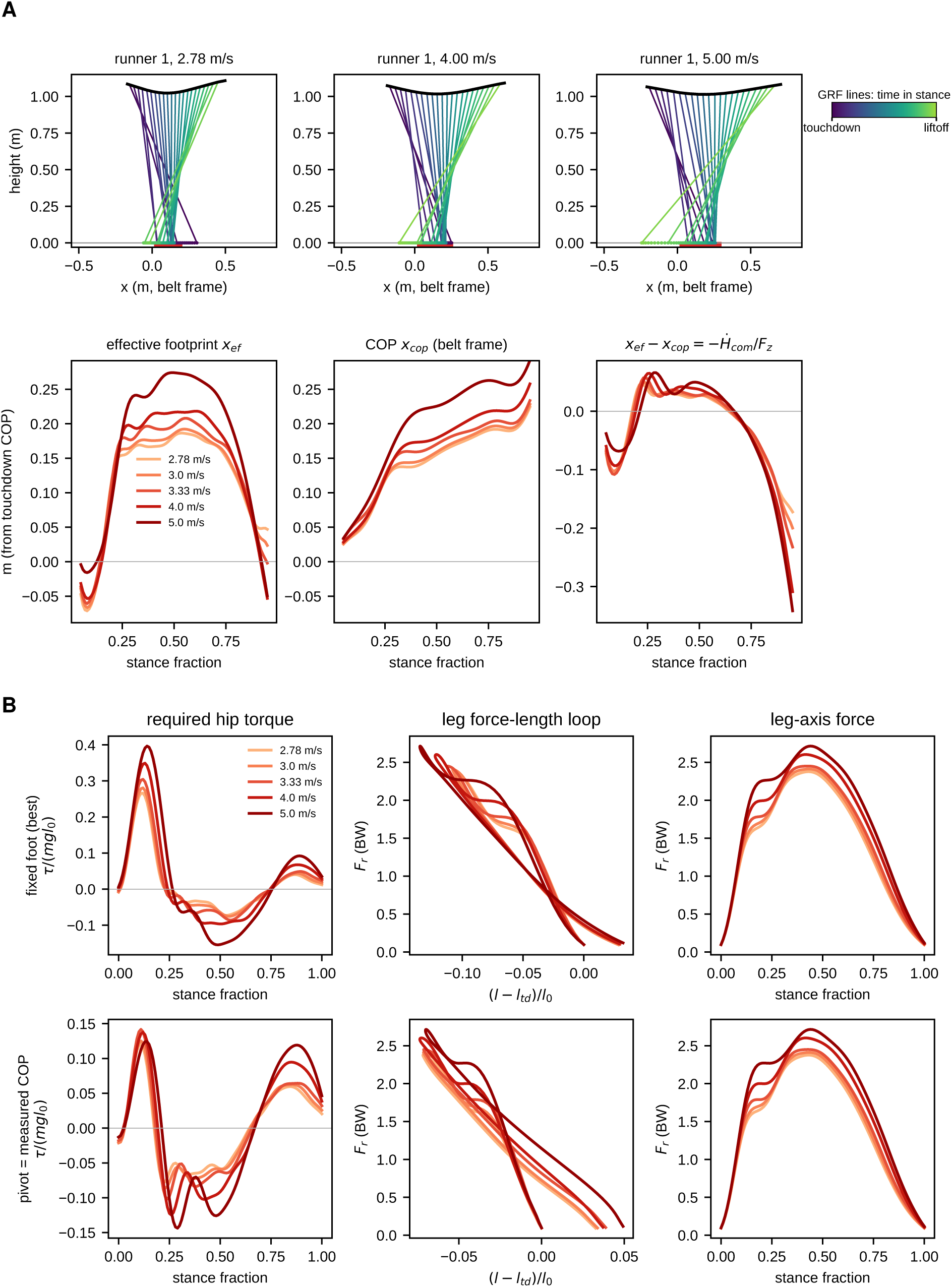
Measured force-line geometry and the hip torque a template needs. (A) Top: GRF lines through the COM for runner 1 of Van Hooren and Meijer (2024) at three speeds, coloured from touchdown to liftoff as in Fig. 3, with the effective footprints as dots on the ground (red bar, COP travel in the belt frame). Bottom: means over 19 runners of the effective footprint, the COP and their difference *x_ef_ −x_cop_* = −*Ḣ /F_z_* (5–95% of stance, *F_z_ >* 0.2 BW). (B) Template inversion of the condition means: hip torque, leg-axis force–length loop and leg-axis force a point-mass template must produce, on the best fixed foot (top) and with the pivot at the measured COP (bottom). In the speed-resolved panels, colours

### The complexity ladder on one dataset

On the 95 conditions of Van Hooren and Meijer (2024), the present pipeline reproduced the 2012 errors on the digitized curves within 2% (Table S10), and the 34 templates separated into two groups (Fig. 5). Every template without leg damping — SLIP, the power-law spring, the moving foot, the point-mass VPP, the clock-torqued SLIP and the hip spring–damper without leg damping — stayed at 0.35–0.36 BW, close to SLIP (0.363 BW); the two-segment knee leg was worse (0.517 BW). Templates with leg damping and a non-radial element reached 0.16–0.23 BW. The best template was TD-SLIP-2d (RMSE 0.160 BW; fore-aft 0.080, vertical 0.138), ahead of all others by at least 471 summed BIC units and BIC-best in 51 of 95 conditions (Table 1, Table S3). It also had the lowest median test error in all three cross-validation schemes: 0.165 BW for a new speed, 0.183 BW for a new runner, and 0.199 BW with one parameter set for all runners (next best 0.177, 0.196 and 0.202 BW; Table S4 gives means).

**Figure 5:**
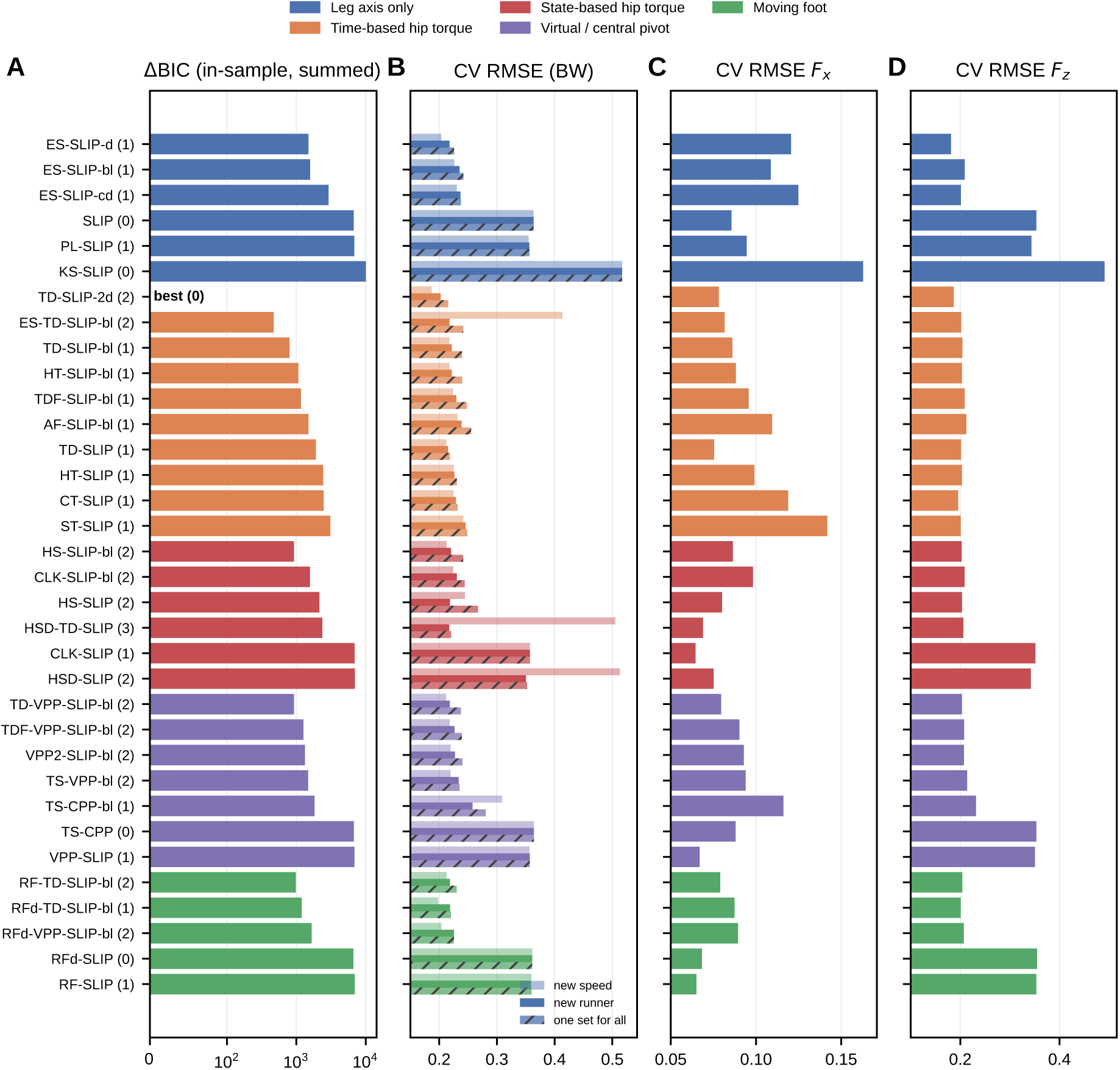
Benchmark on one dataset (95 conditions of Van Hooren and Meijer 2024), grouped by family. (A) BIC excess summed over conditions relative to the best template (symmetric-log axis; the best, TD-SLIP-2d, is marked). (B) Cross-validated test RMSE for a new speed (light), a new runner (solid) and one parameter set for all runners (hatched). (C,D) Test RMSE of the fore-aft and vertical force for a new runner. Numbers in brackets give the number of shape parameters.

**Table 1:** Key templates. *p*, shape parameters fitted to the GRF. RMSE (BW) of free fits on Van Hooren and Meijer (2024) for the combined, fore-aft (*x*) and vertical (*z*) force; *F_x_* floor, best fore-aft error when the fore-aft force is fitted alone. Remaining columns: mean BIC excess per condition relative to the best template of each dataset (bold, best) and its maximum over datasets. VH, Van Hooren; FU, Fukuchi; CA, Carter; BK, Baker treadmill; BKO, Baker overground; RG, Riglet; max, maximum over the human datasets; GF, guinea fowl (Daley and Birn-Jeffery, 2026b); GH and GL, guinea fowl on high-and low-friction ground (Goldsmith et al., 2026). All 34 templates in Tables S3–S7.

|  |  | Van Hooren, RMSE (BW) |  |  |  | BIC excess per condition |  |  |  |  |  |  |  |  |  |
|---|---|---|---|---|---|---|---|---|---|---|---|---|---|---|---|
|  |  |  |  |  |  | humans |  |  |  |  |  |  | guinea fowl |  |  |
| Template | *p* | all | *x* | *z* | *F<sub>x</sub>* floor | VH | FU | CA | BK | BKO | RG | max | GF | GH | GL |
| *Leg axis only* |  |  |  |  |  |  |  |  |  |  |  |  |  |  |  |
| SLIP | 0 | 0.363 | 0.086 | 0.353 | 0.086 | 71 | 73 | 64 | 69 | 72 | 60 | 73 | 59 | 70 | 57 |
| ES-SLIP-d | 1 | 0.194 | 0.118 | 0.154 | 0.085 | 16 | 13 | 9 | 32 | 30 | 14 | 32 | 26 | 46 | 38 |
| *Time-based hip torque* |  |  |  |  |  |  |  |  |  |  |  |  |  |  |  |
| TD-SLIP | 1 | 0.202 | 0.075 | 0.187 | 0.073 | 20 | 22 | 17 | 45 | 44 | **0** | 45 | 34 | 51 | 37 |
| TD-SLIP-bl | 1 | 0.186 | 0.087 | 0.165 | 0.080 | 8 | 7 | 5 | 5 | 3 | 27 | 27 | 32 | 47 | 40 |
| TD-SLIP-2d | 2 | 0.160 | 0.080 | 0.138 | 0.073 | **0** | **0** | **0** | 9 | 7 | 1 | 9 | 30 | 47 | 37 |
| ES-TD-SLIP-bl | 2 | 0.169 | 0.078 | 0.150 | 0.054 | 5 | 7 | 6 | 8 | 6 | 24 | 24 | 34 | 50 | 43 |
| *State-based hip torque* |  |  |  |  |  |  |  |  |  |  |  |  |  |  |  |
| HS-SLIP-bl | 2 | 0.180 | 0.081 | 0.161 | 0.066 | 10 | 9 | 8 | 9 | 7 | 29 | 29 | 19 | 23 | 18 |
| CLK-SLIP-bl | 2 | 0.194 | 0.095 | 0.169 | 0.071 | 17 | 14 | 12 | 11 | 10 | 34 | 34 | 30 | 37 | 33 |
| HSD-TD-SLIP | 3 | 0.195 | 0.058 | 0.186 | 0.056 | 25 | 31 | 25 | 56 | 55 | 8 | 56 | 32 | 38 | 26 |
| *Virtual / central pivot* |  |  |  |  |  |  |  |  |  |  |  |  |  |  |  |
| VPP-SLIP | 1 | 0.356 | 0.063 | 0.350 | 0.061 | 72 | 75 | 67 | 72 | 74 | 62 | 75 | 50 | 48 | 38 |
| VPP2-SLIP-bl | 2 | 0.188 | 0.087 | 0.167 | 0.061 | 14 | 11 | 9 | 7 | 3 | 32 | 32 | 1 | 0 | 0 |
| TD-VPP-SLIP-bl | 2 | 0.179 | 0.076 | 0.162 | 0.061 | 10 | 7 | 7 | 7 | 4 | 29 | 29 | 2 | 2 | 1 |
| TS-CPP | 0 | 0.364 | 0.088 | 0.353 | 0.088 | 71 | 73 | 64 | 70 | 72 | 60 | 73 | 60 | 76 | 59 |
| TS-CPP-bl | 1 | 0.208 | 0.111 | 0.176 | 0.088 | 19 | 17 | 11 | 15 | 12 | 33 | 33 | 50 | 64 | 50 |
| *Moving foot* |  |  |  |  |  |  |  |  |  |  |  |  |  |  |  |
| RF-TD-SLIP-bl | 2 | 0.181 | 0.075 | 0.165 | 0.060 | 10 | 6 | 8 | **0** | **0** | 30 | 30 | 8 | 15 | 15 |
| RFd-VPP-SLIP-bl | 2 | 0.195 | 0.086 | 0.176 | 0.060 | 17 | 12 | 11 | 7 | 3 | 36 | 36 | **0** | 4 | 4 |

Fitting each force component alone showed what each element fixes (Table 1, Table S5). Leg-axis templates barely improved the fore-aft force: their best fore-aft-only errors were 0.072 BW (power-law spring) and 0.085–0.088 BW (damped and switching legs), against 0.086 BW for SLIP, because a radial force is tied to the leg angle. The best hip-torque and pivot templates lowered it to 0.054–0.061 BW (the TD-SLIP ramp alone: 0.073 BW). Conversely, leg damping is what improves the vertical force (best vertical-only errors 0.139–0.154 BW, against 0.353 for SLIP). A leg-axis template with linear damping and a force-continuous stiffness switch (ES-SLIP-d) had the best vertical error of all one-parameter templates but a fore-aft error of 0.12 BW.

### State-based hip laws and pivots

A hip torsional spring with its rest angle near the liftoff angle fitted as well as the time-based ramp, with which it is nearly equivalent, but did not earn its extra parameter (HS-SLIP-bl: 925 BIC units behind TD-SLIP-2d, against 798 for the one-parameter ramp with bilinear damping); the clock-torqued SLIP was slightly worse (CLK-SLIP-bl, 1570). Adding a hip damper to the hip spring of a damped TD-SLIP (HSD-TD-SLIP, three parameters) gave the lowest fore-aft error of all templates (0.058 BW) and the only effective footprint that swings back late in stance as the data do (Fig. 6C), but its vertical force was limited by linear damping and it did not earn its parameters. The trunk template with the CPP reproduced SLIP to within 0.001 BW, as expected because routing the force through the COM makes it central; with damping it could close the energy balance only through a forward trunk lean of 26^◦^ (median). VPP templates improved the fore-aft force but placed the fitted pivot above the COM (median 0.16–0.25 *l*_0_), opposite to the measured pivot 0.27–0.31 m below the COM (Fig. 4A), and none earned its parameter.

**Figure 6:**
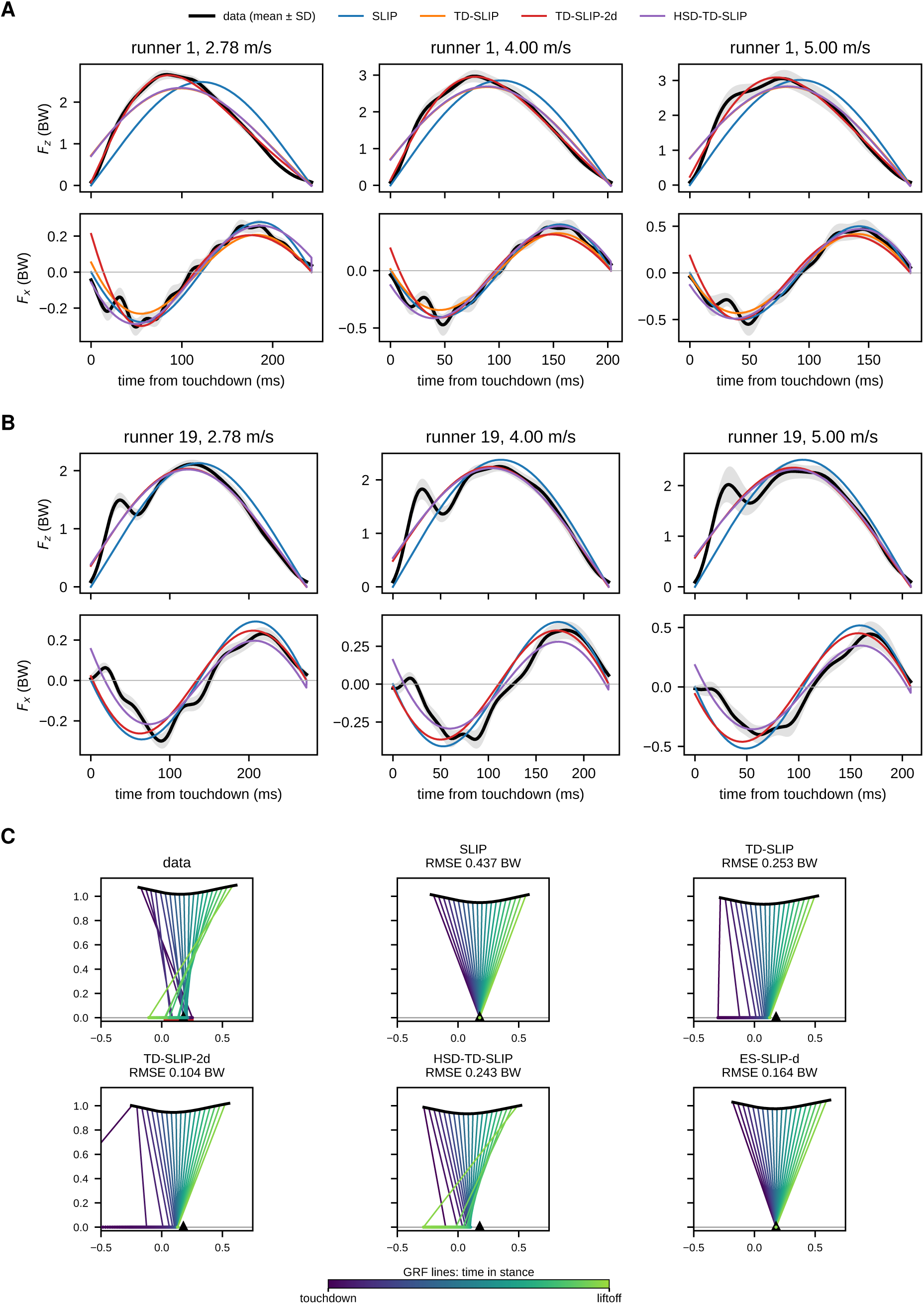
Fitted templates against measured GRF. (A) Runner 1 of Van Hooren and Meijer (2024) (early vertical peak) and (B) runner 19 (heel strike with a distinct impact peak) at three speeds: condition-mean vertical (top) and fore-aft (bottom) force (±s.d. over steps) with SLIP, TD-SLIP, TD-SLIP-2d and the hip spring–damper TD-SLIP. (C) Effective footprints of runner 1 at 4 m s^−1^: GRF lines through the COM coloured from touchdown to liftoff as in Fig. 3, for the data (red bar, COP travel; triangle, best fixed foot) and the fitted templates. Only the hip spring–damper template reproduces the late-stance swing of the footprints.

### Replication across six human datasets

Across datasets (Fig. 7, Table 1, Tables S6–S7) the floor and the top of the ladder were the same. SLIP, the CPP and all undamped templates were 56–78 BIC per condition behind the best template in every dataset (the two-segment leg 101–150). TD-SLIP-2d was the best template on Van Hooren, Fukuchi (lead over the next template 576 BIC units; 39% of conditions) and Carter (lead 462; 41%), and within 1.5, 7.1 and 9.3 BIC per condition of the best template on Riglet, Baker overground and Baker treadmill — the smallest worst case of all 34 templates. Its in-sample error was 0.15–0.18 BW in all datasets (SLIP: 0.35–0.37). With one parameter set for all conditions it had the lowest median test error on Van Hooren (0.199 BW), Carter (0.201) and Riglet (0.174) and was within about 0.02 BW of the best elsewhere (Table S7). The AIC agreed with the BIC on the best template in five of the six datasets (rank correlation over all templates 0.978–0.994). Where the two differed, the AIC preferred the template with more parameters: TD-SLIP-2d over the one-parameter TD-SLIP on Riglet, and two-parameter pivot templates entered its top three on Van Hooren, Fukuchi and both Baker datasets.

**Figure 7:**
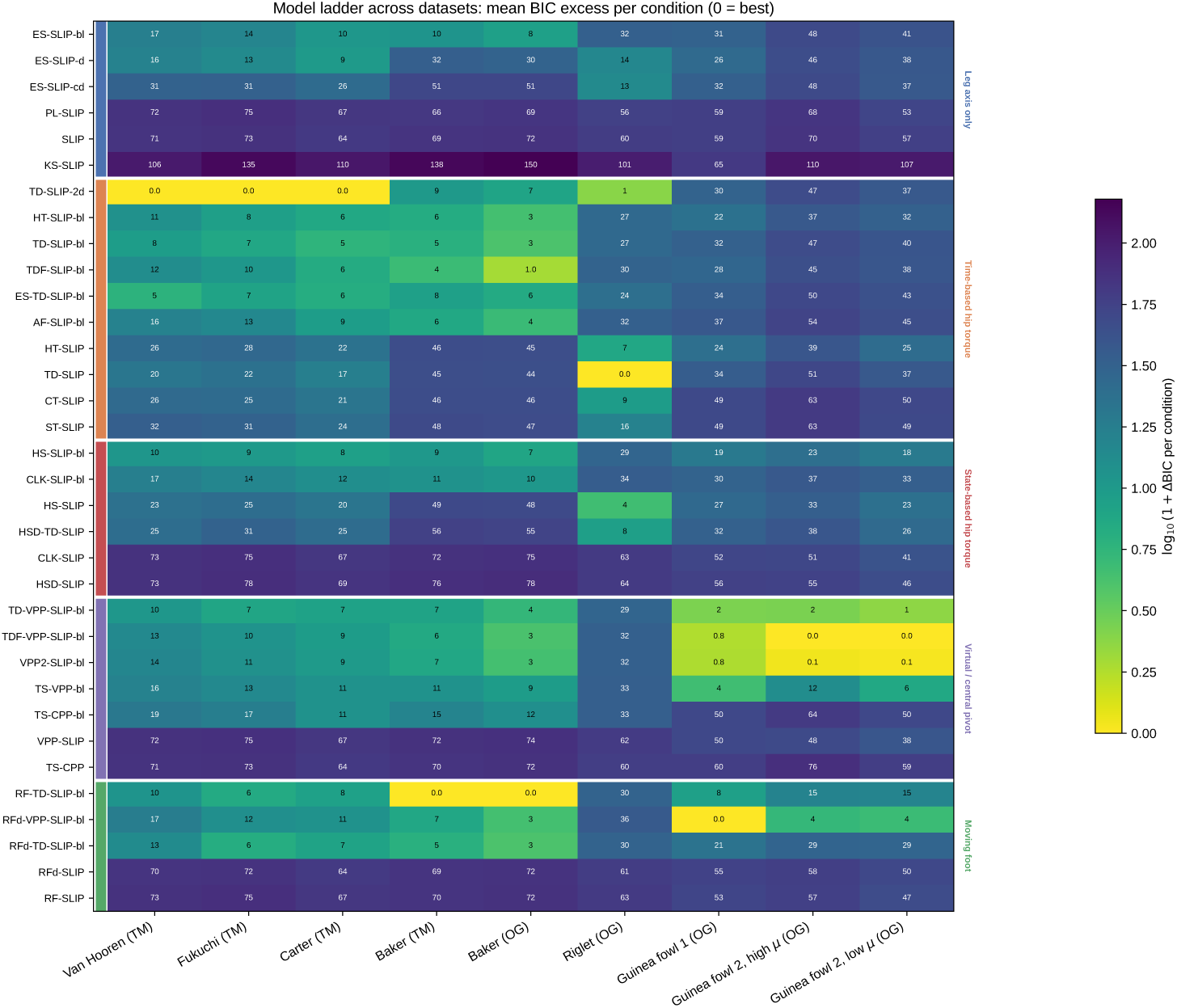
Replication across six human and two guinea-fowl datasets. Mean BIC excess per condition relative to the best template of each dataset (0 = best; colour on a log scale), grouped by family. TM, treadmill; OG, overground; guinea fowl 1, Daley and Birn-Jeffery (2026b); guinea fowl 2, Goldsmith et al. (2026) on high- and low-friction (*µ*) ground. Datasets in Table S1; numbers in Table S6.

### A running bird needs the same elements in another form

In guinea fowl the floor of the ladder was the same as in humans (Fig. 7, Table 1): templates without leg damping were 50–60 BIC per condition behind the best (two-segment leg 65; SLIP: 0.47 BW; best undamped template 0.37 BW), and a damped leg-axis template (ES-SLIP-d) left a fore-aft error of 0.20 BW. The top, however, differed. All four point-mass templates that combine bilinear damping with a virtual pivot were within 1.7 BIC per condition of the best (RFd-VPP-SLIP-bl, whose foot follows the measured stance foot: 0.127 BW; fore-aft 0.053), and the trunk template with a trunk-fixed pivot followed (3.7); the best hip-torque template moved its foot (RF-TD-SLIP-bl, 7.9), and TD-SLIP-2d was 30.4 BIC per condition behind. The fitted pivots lay 0.85–1.15 *l*_0_ above the COM (medians of the four point-mass pivot templates), and the measured force lines pointed the same way: their least-squares intersection, with the stance foot as the COP, lay 0.53 *l*_0_ above the COM (interquartile range 0.46–0.60), whereas in humans it lay 0.25 *l*_0_ below it. Cross-validation agreed: with one parameter set for all conditions, tested on a held-out bird, the pivot templates reached median errors of 0.142–0.144 BW, against 0.187 for RF-TD-SLIP-bl and 0.277 for TD-SLIP-2d.

The second guinea-fowl dataset, with a measured COP, gave the same answer on both surfaces (Fig. 7, Table 1). The point-mass pivot templates with bilinear damping were again best, within 1.7 BIC per condition of each other (best errors 0.144 and 0.139 BW on high and low friction; fore-aft 0.043 and 0.037 BW), with fitted pivots 0.85–1.30 *l*_0_ above the COM. The best hip-torque template (RF-TD-SLIP-bl) was 15 BIC per condition behind on both surfaces and TD-SLIP-2d 47 and 37; templates without leg damping were 38 or more behind, and the damped leg-axis template again left a large fore-aft error (0.21 and 0.17 BW). The measured force lines, now through the measured COP, met 0.58 *l*_0_ (high friction; interquartile range 0.51–0.73) and 0.67 *l*_0_ (low friction; 0.55–0.71) above the COM. With one parameter set for all conditions and a held-out bird, the moving-foot pivot template reached median errors of 0.093 and 0.112 BW, against 0.161 and 0.120 for RF-TD-SLIP-bl and 0.276 and 0.194 for TD-SLIP-2d. The pivot height depends on where the COM is: with the exported COM height (about 0.08 m higher), the force lines still met above the COM (0.14 and 0.18 *l*_0_), the pivot templates were still BIC-best on both surfaces with fitted pivots 0.25–0.70 *l*_0_ above the COM, but the margin to the hip-torque templates shrank to 6–12 BIC per condition and to the damped leg-axis template to 7–18 (Table S12).

Slippery ground did not change the template. In the ten bird–speed conditions present on both surfaces, peak vertical force (1.76 and 1.75 BW) and stance time (0.165 and 0.158 s) were unchanged, and the fore-aft forces were, if anything, larger on the slippery surface (peak propulsion 0.21 versus 0.23 BW, Wilcoxon *P* = 0.02; peak braking 0.27 versus 0.28 BW, *P* = 0.08). The force-line intersection (0.57 versus 0.63 *l*_0_, *P* = 0.38), the fitted pivots (median shift less than one grid step, *P* 0.06) and the damping constants did not change systematically, and the ranking was the same on both surfaces (Table S13). These are steady aerial steps that did not slip, so they show what a bird does when it keeps running on a low-friction surface, not how it recovers from a slip.

The preferred damping law, however, differed. On Riglet et al. (2024) (overground, distinct heel strikes) the original TD-SLIP with *linear* damping was BIC-best in 37 of 57 conditions, and bilinear-only templates were 24–36 BIC per condition behind. On Baker et al. (2025) the order was reversed: bilinear-damping templates led and linear-damping TD-SLIP was 44–45 BIC per condition behind; the best template there moved its foot at a fitted speed (RF-TD-SLIP-bl), consistent with the absence of COP data in that dataset. The other three datasets lay in between, where the two-term law won.

### What the damping terms encode

Three analyses addressed the damping law (Fig. 8). First, in TD-SLIP-2d fitted per runner, the bilinear constant fell (*r* = −0.88) and the linear constant rose (*r* = +0.58) with the time of the vertical force peak (Fig. 8B): early-peak runners were bilinear, runners with a distinct impact peak and a later main peak were linear. Second, a foot mass with a soft contact on a bilinear core (IM-TD-SLIP-bl) matched the error of TD-SLIP-2d at 4 m s^−1^ (0.169 vs 0.170 BW) and had the lowest vertical error in the first 15% of stance (0.231 vs 0.270 BW), with a plausible foot touchdown speed of about 0.9 m s^−1^ (Fig. 8A, Table S8); it did not reach the full heel-strike peak and produced too large an early positive fore-aft force. Third, removing the impact transient by a 15 Hz low-pass filter did not change the ranking: TD-SLIP-2d remained first (11 of 19 conditions, lead 195 BIC units) and bilinear-only TD-SLIP stayed 380 BIC units behind (Fig. 8C). The dataset differences are therefore not a bandwidth effect. Neither were they explained by foot-strike category: in Fukuchi et al. (2017), rearfoot and fore/midfoot strikers did not differ in fitted damping constants (Mann–Whitney *P* = 0.17 and 0.34) and preferred bilinear over linear damping equally often (70% and 68% of conditions).

**Figure 8:**
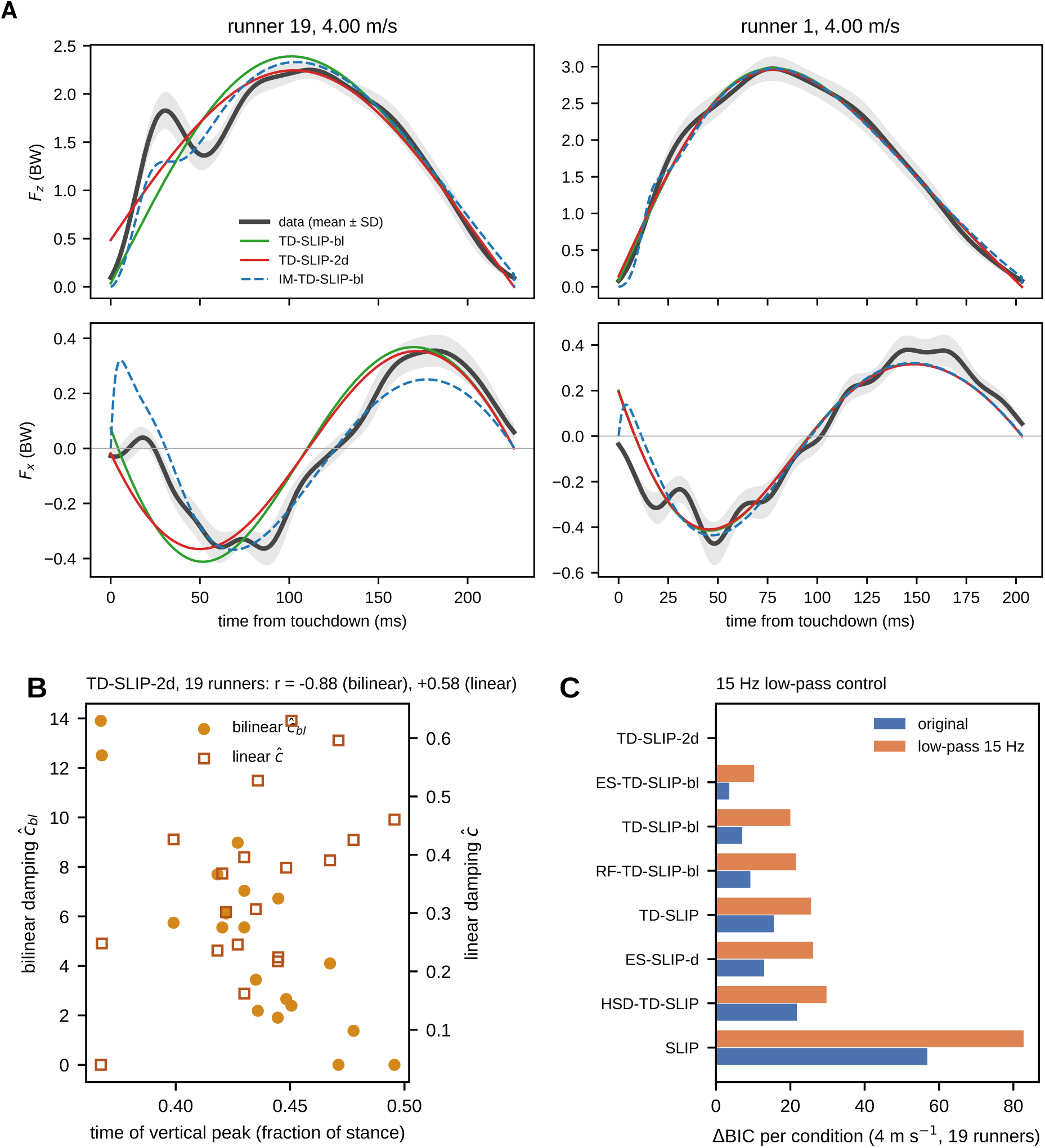
What the damping terms encode. (A) Foot-mass template on a bilinear core (IM-TD-SLIP-bl) compared with the bilinear TD-SLIP and TD-SLIP-2d for a heel striker (runner 19) and an early-peak runner (runner 1) at 4 m s^−1^. (B) Fitted bilinear (filled) and linear (open) damping of TD-SLIP-2d against the time of the vertical force peak, one point per runner (means over speeds). (C) BIC excess per condition of eight templates on the 19 conditions at 4 m s^−1^, original and after a 15 Hz low-pass filter that removes the impact transient.

The parameters varied between runners more than the gait they were fitted to. On Van Hooren and Meijer (2024), the measured peak vertical force, dimensionless stance time and touchdown angle had coefficients of variation of 7–13% across the 19 runners at each speed, whereas the fitted parameters of TD-SLIP-2d varied by 17–26% (stiffness), 20–29% (torque amplitude), 45–49% (linear damping) and 66–79% (bilinear damping). The two damping constants traded off against each other (*r* = −0.68 to −0.82 across runners). One parameter set for all runners raised the error from 0.160 BW (per-condition fits) to a median of 0.199 BW on held-out runners. The lag between the fore-aft braking–propulsion transition and the vertical force peak, a timing index of spring–mass similarity (Burns et al., 2023), had medians of 17–21 ms across speeds but was not related to the fitted torque amplitude across runners (Spearman *ρ* = 0.07–0.17, *P* 0.1, TD-SLIP and TD-SLIP-2d).

### Treadmill versus overground running

For the same 15 runners (Baker et al., 2025), the ranking of all 34 templates was the same overground and on the treadmill (Spearman *ρ* = 0.998; Fig. 9A). The parameters changed consistently (Fig. 9B, Table S9): overground, the fitted dimensionless stiffness was lower (median paired difference −8%; *k*^^^ 16.9 to 15.4, *P* = 0.002, in TD-SLIP, its bilinear and its two-term variant), stance was 12 ms longer (*P* = 0.011), the touchdown angle was slightly flatter (+0.007 to +0.014 rad, *P* = 0.03–0.06) and the vertical peak about 0.1 BW lower (*P* = 0.055), while the torque amplitude and damping constants did not change. Across datasets at matched speed (Riglet against the three treadmill datasets with COP), overground conditions had more linear and much less bilinear damping (*c*^ +0.39, 95% CI 0.31–0.46; *c*^*_bl_* −4.5, −5.3 to −3.5) and a 0.24 BW lower vertical peak, but no difference in stiffness, torque amplitude or touchdown angle (Fig. 9C–F). Because the same runners show no change in the damping mix, this cross-dataset difference reflects the runner samples rather than the setting.

**Figure 9:**
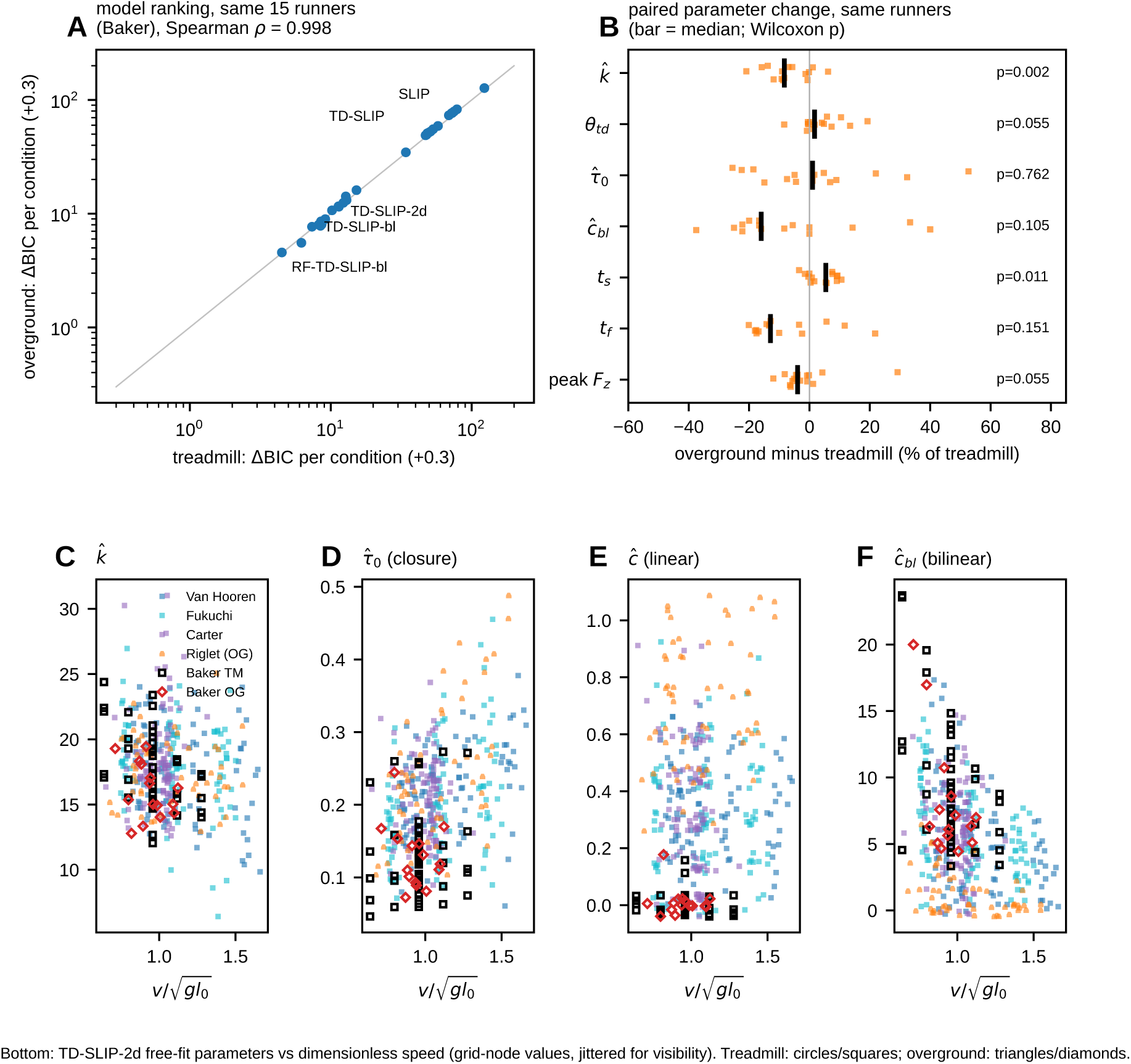
Treadmill versus overground running. (A) BIC excess per condition of all 34 templates for the same 15 runners overground against the treadmill (Baker et al., 2025). (B) Paired overground-minus-treadmill changes of fitted parameters and gait variables (dots, runners; bar, median; Wilcoxon signed-rank *P*). (C–F) TD-SLIP-2d parameters against dimensionless speed in all datasets (grid-node values, jittered for visibility): stiffness *k*^^^, torque amplitude *τ*^_0_, linear and bilinear damping.

### The hip torque a template needs

Template inversion on the condition means (Fig. 4B) showed what a point-mass template on a fixed foot must produce to match the data exactly. The best fixed foot lay 0.15–0.21 m ahead of the touchdown COP. The required hip torque was tri-phasic: a positive pulse peaking near 12% of stance (0.27–0.40 *mgl*_0_, increasing with speed), a negative phase in mid-stance and a small positive bump before toe-off. The leg-axis force did net negative work (0.010 to 0.018 *mgl*_0_ per step; its negative part alone 0.15 to 0.22 *mgl*_0_), and the hip torque supplied the same amount. The TD-SLIP ramp captures the first phase and this energy exchange; the hip spring–damper captures all three phases. The fitted templates did much more hip work than this minimum. In TD-SLIP-2d the hip torque supplied, and the leg damping dissipated, 0.067–0.135 *mgl*_0_ per step (medians over runners, increasing with speed), about seven times the work found by inversion. Its foot lay 6–7 cm ahead of the best fixed foot, and because *x_ef_ x_f_* = *τ/F_z_*, a foot further ahead of the same force lines needs a larger forward torque.

## Discussion

### What is necessary

Across six human datasets and a bird, two elements were necessary and, together, nearly sufficient. A non-central element — a hip torque, a pivot or a moving contact — is needed for the fore-aft force, because a force along the leg leaves the fore-aft error near the SLIP value whatever the leg law. Leg damping is needed for the vertical force: no undamped template brought its vertical error below 0.30 BW, whatever its hip law. The templates with the fewest parameters that include both belong to the TD-SLIP family proposed in 2012 (Ankaralı et al., 2012): its two-parameter form with linear plus bilinear damping is within 9.3 BIC per condition of the best template in every human dataset and is the most reliable single choice for human running, and the original one-parameter TD-SLIP is the best template where impact transients are pronounced. This is a nested test of necessity, of the kind that showed that cockroach wall following needs derivative feedback (Cowan et al., 2006): an element is necessary when no template without it, whatever its other terms, reaches the error of templates with it. Nothing with more than two shape parameters was consistently worth its cost, and choosing the AIC instead of the BIC would have added at most one parameter to the selected template without changing which elements are necessary. These conclusions are stable across laboratories, runner samples, treadmill and overground running, force bandwidth, and two definitions of generalisation.

In the terms of Full and Koditschek (1999), the benchmark selects the template; it does not select the anchor. The steps towards an anchor that the ladder contains — a two-segment leg with a knee, a trunk with a hip joint, a foot mass with a soft contact — did not improve the account of the GRF, with one exception: the foot mass could replace the linear damper. Added morphology therefore bears on how the template is realised, not on which template is realised, and that is a question for joint-level data.

The bird shows where morphology does enter. Guinea fowl needed the same two elements as humans, but their non-central force took the form of a virtual pivot above the COM rather than a ramp of hip torque. A pivot above the COM is the arrangement that stabilises trunk pitch in walking and running models (Maus et al., 2010), and a bird’s long, nearly horizontal trunk with the COM ahead of the hips makes pitch control a larger part of the force pattern than in an upright human. The species thus differ in the anchor’s constraint, not in the template’s elements: both pay dissipation along the leg back through a moment at the hip. The pivot was not an artefact of one laboratory’s processing: it reappeared in a second set of birds with a measured COP rather than a foot marker, and on slippery as on grippy ground. Its height, however, rests on the COM estimate. Raising the COM by 0.08 m, as in the exported data, lowered the pivot to 0.25–0.70 *l*_0_ above the COM and narrowed the lead over hip-torque templates, so in birds the sign of the pivot (above, not below, the COM) is firm, whereas its height needs a COM that is measured rather than reconstructed.

Low friction did not move the bird to another template. On the slippery surface the birds that kept running in aerial steps produced the same vertical force and, if anything, slightly larger fore-aft forces, with the same ranking and no systematic change of pivot or damping. A low friction coefficient bounds the ratio of fore-aft to vertical force at the foot; in these steady steps that bound was evidently not reached. The finding that gave Goldsmith et al. (2026) their title, that faster birds fell more often on slippery ground, points to speed rather than the stance force pattern as what the birds manage there. The template describes what a stance does once the foot holds; how a bird recovers when it does not lies outside it.

A likely root of this difference is how body mass is divided between the trunk and the limbs. A human leg carries about 16% of body mass and an arm about 5% (Winter, 2009), whereas a guinea-fowl leg carries about 13% (Rubenson and Marsh, 2009): the body-to-leg mass ratio is about 6 in humans and 8 in the bird, and the swinging limbs together hold about 40% of a human’s mass but about a quarter of the bird’s. Because *x_ef_ −x_cop_* = −*Ḣ _com_/F_z_*, a GRF line that misses the COM changes the whole-body angular momentum, which the body must absorb and return within the stride. Heavy, counter-swinging legs and arms can carry that angular momentum, so a human can let the force pass below the COM; in a bird whose mass is concentrated in the trunk, the same moment would pitch the trunk, and a force directed through a point above the COM keeps the moment small and restoring (Maus et al., 2010). On this view the choice between a time-based hip torque and a virtual pivot is set by the anchor’s mass distribution, and running animals with light limbs and heavy trunks should favour pivot templates.

The two comparisons that matter most run in opposite directions. The time symmetry of SLIP’s stance is not symmetric enough: the undamped templates were 56–78 BIC per condition behind the best in every human dataset and 38 or more in the birds, so the asymmetry of the measured force must be modelled. The difference between treadmill and overground running is the reverse case, as for left–right asymmetry in walking (Ankaralı et al., 2015): the same runners had a measurably lower stiffness and longer stance overground, but the ranking of all 34 templates was unchanged (*ρ* = 0.998), so for choosing a template treadmill running is overground enough.

### Biological interpretation

In template terms, running is a lossy spring whose losses are paid back by hip work. The template inversion makes this explicit: about 0.01–0.02 *mgl*_0_ per step is dissipated along the leg and supplied by the hip, and the required hip torque is largest early in stance, when the leg is braking. The damping terms map onto two physical sources of loss. The linear term, with its force jump at touchdown, stands in for the collision of the foot and shank with the ground: a 6% foot mass with a soft contact on a bilinear core reproduces its effect and the early vertical force. The bilinear term, which grows with compression, encodes the skew of the main vertical peak and dominates in runners whose force peaks early. That the mix tracked peak timing but not foot-strike category suggests that peak timing reflects more than the initial contact, for example knee and ankle compliance during weight acceptance.

Dissipating energy only to resupply it looks wasteful, as do the braking and propulsive impulses that cancel in every stance. Mutually opposing forces need not be wasteful: in the glass knifefish, two counter-propagating fin waves whose thrusts cancel add passive damping and make small manoeuvres cheap, removing the trade-off between stability and manoeuvrability (Sefati et al., 2013). Leg damping paired with hip actuation is an arrangement of the same kind. Energy could in principle be returned along the leg instead: with virtual damping tuned by an actuator in series with the leg spring, a robot leg adds and removes energy through the leg axis alone (Seçer and Saranlı, 2014, 2018). That suffices for a template whose task is the COM trajectory, but not for the human or bird GRF: the leg-axis template that closes its energy balance through a stiffness switch at mid-stance (ES-SLIP-d) fitted the vertical force and left a fore-aft error of 0.12 BW in humans and 0.20 BW in guinea fowl. Running forces require at least part of the energy lost along the leg to be returned through a force off the leg axis. In a torque-actuated dissipative spring–mass hopper it yields stride-to-stride energy regulation and self-stability (Ankaralı and Saranlı, 2010), and the hip torque is a handle through which a controller can change speed. Full and Koditschek (1999) proposed that in rapid running a tuned mechanical system rejects disturbances passively and simplifies control; ATRIAS was built on the same premise, with tuned compliance providing stabilising passive dynamics that the controller exploits (Hubicki et al., 2018); and in a hopping robot tunable leg damping improved recovery from perturbations by up to 170% at a 27% higher energetic cost (Mo et al., 2023): damping trades efficiency for robustness. The benchmark shows that running forces require the pair; whether runners use it for stability is a question for perturbation experiments.

The GRF fixes the template but not its anchoring: it fixes the sum of the leg and hip actions but not how it is split between them. The same forces were reproduced with about 0.01–0.02 *mgl*_0_ of hip work per step on the best fixed foot and with about seven times as much by the fitted TD-SLIP-2d, whose foot lay a few centimetres further forward, and time-based and state-based hip laws fitted equally well. Closed-loop behaviour constrains the loop more than its parts: in refuge tracking by electric fish, a broad family of controllers is consistent with the same behaviour (Uyanik et al., 2020). Template parameters are therefore effective properties of the closed loop, body mechanics and neural feedback together. Legged robotics arrived at the same view from the other side: a series-actuated leg can be driven to behave as a damped SLIP whose damping coefficient is set by the controller, and tuning that virtual damping regulated running with less peak actuator power and less wasted negative work than modulating leg stiffness (Seçer and Saranlı, 2014, 2018). A fitted damping constant is likewise a virtual damping: the net effect of passive tissue losses and active negative work at the joints, which a runner can in principle adjust. Small changes in neural drive can shift the effective mechanical properties of a body (Tytell et al., 2011), and the same muscle activation produces different motion when body stiffness changes, with negative muscle work emerging from the interaction of body and environment (Tytell et al., 2010). The treadmill–overground comparison fits this reading: the effective stiffness changed with the surface while the torque and damping did not.

Between runners, the template parameters had coefficients of variation two to nine times those of the gait observables, and the two damping constants traded off. Part of this is degeneracy, since different mixes of linear and bilinear damping give similar forces, and part is real, since the mix followed the timing of the force peak. As in electric fish, where consistent tracking emerges from variable plants (Uyanik et al., 2020), what is common to runners is the structure of the template rather than its parameter values, which is why templates should be compared on held-out runners.

The fact that hip-based and pivot-based templates ranked similarly, and that state-based hip springs fitted as well as the time-based ramp with a rest angle near liftoff, shows that the GRF alone cannot distinguish time-based from state-based hip control at this level of detail. Likewise, the CPP, which simplifies trunk control and makes the trunk dynamics decouple (Sever et al., 2026), is a posture law rather than a GRF template: by construction it produces a central force and therefore SLIP’s GRF. The measured pivot below the COM is a statement about force lines through the moving COP (Drama et al., 2020); a point-mass template with a fixed foot reproduces the same forces with a hip torque and places any fitted pivot above the COM. In walking, too, the VPP is always observed, yet simulations suggest that it may not be necessary for stability (Schreff et al., 2023); a pivot is one way to obtain a non-central force, not a requirement of the data.

The template’s foot is not the anatomical contact point. The ramp template driven by the measured COP (RFd-TD-SLIP-bl) fitted worse than the same template on a fixed foot in three of the four datasets with COP data (Table S6), and the best fixed foot lay near the mid-stance COP. The effective footprint, not the COP, is the quantity the template reproduces, which is why the footprint picture is the natural lens for comparing templates and data.

### Implications for legged robots

Because templates carry biological insight into robot design and control, the benchmark bears on machines in several ways. A robot that embeds SLIP reproduces the COM motion of running but not its forces; where animal-like forces matter, for example for loading of the structure or for devices that interact with people, the template it embeds should include leg damping and a force off the leg axis. Neither element needs dedicated hardware. Damping can be produced by an actuator in series with the leg spring (Seçer and Saranlı, 2014, 2018), and a hip torque or a virtual pivot by a hip actuator (Sharbafi et al., 2013); because the fitted parameters are effective properties of the closed loop, a machine can match them by control. The time-based family that the human data select is also one that robots already run on: a clock-driven hip torque on a SLIP leg, close to the TD-SLIP ramp in our benchmark (CLK-SLIP-bl), carries a quadruped from walking to running trot (Akbaba et al., 2025).

Two differences between animals and machines stand out. First, the routes by which energy is returned differ. Robot controllers can regulate energy efficiently along the leg (Seçer and Saranlı, 2018), whereas human and bird forces require that part of it comes back through a non-radial force. A dissipative leg paired with a hip torque gives stride-to-stride energy regulation and self-stability (Ankaralı and Saranlı, 2010), so the animal route may buy robustness and a handle on speed rather than efficiency, as tunable damping does in a hopping robot (Mo et al., 2023). Second, the roles of the off-axis force differ. In hierarchical template-based control the COM template and the body’s orientation are handled by separate loops (Ankaralı and Saranlı, 2011; De et al., 2022), so the force off the leg axis belongs to the posture loop. The animals do not separate them in this way: in humans the off-axis force, as a hip torque, returns the energy that the leg dissipates, and in guinea fowl it takes the form of a pivot that also governs trunk pitch. A robot template that reproduces animal forces would therefore couple the energy balance of the COM template to the posture channel. Which form that channel should take depends on the body plan. Most legged robots are built like the bird rather than the human, with light legs and the mass in the body (Hubicki et al., 2016); by the mass-distribution argument above they should be better served by a virtual pivot than by a hip-torque program, a hypothesis that is easier to test on a machine than on an animal. Finally, the benchmark can be run on a robot’s own GRF to measure how closely the machine embeds the template it was designed for. This extends to the whole template ladder the identification of a monopod’s SLIP parameters that preceded validation of the analytic stance predictor on hardware (Uyanık et al., 2015).

### Testable predictions

Anchoring the selected template (Full and Koditschek, 1999) gives predictions that can be tested with joint-level data. (i) The net hip moment and power during the first third of stance should scale with the fitted torque amplitude *τ*^_0_ across speeds and runners. Its integral decides how the force is split: the GRF allows between about 0.01–0.02 *mgl*_0_ (best fixed foot) and 0.07–0.14 *mgl*_0_ (fitted TD-SLIP-2d) of hip work per step. Inverse-dynamics hip moments in the Fukuchi and Carter datasets allow this test. (ii) The linear damping constant should scale with the effective mass and stiffness of the foot–ground collision: footwear or surface manipulations that soften the impact should lower *c*^ but not *c*^*_bl_*. (iii) The lower stiffness overground (about 8%) predicts a longer stance and a lower peak force at equal speed and cadence in any overground–treadmill comparison, without a change in hip work. (iv) If the pivot form of the hip moment follows from a horizontal trunk, other running birds and small mammals should also prefer a pivot above the COM, and its height should scale with trunk length; grounded running, which needs a two-leg template, remains untested. (v) If the fitted torque amplitude and damping measure how far a runner departs from SLIP, they should be smaller in runners closer to a symmetric bounce, such as elite distance runners (Burns et al., 2023). Because the transition lag alone did not follow the torque amplitude, such a template-based measure of spring–mass similarity would differ from the timing indices. (vi) If leg damping is a virtual damping set by control, as in series-actuated robot legs (Seçer and Saranlı, 2018), step-by-step fits around an unexpected drop or rise in the ground should show the damping constant changing in the direction that restores the step’s energy, rather than staying at its steady-state value.

### Limitations

Condition means remove step-to-step variability, and stance and flight times are inputs, so the templates predict force shape, not gait timing. Fits other than on Van Hooren are grid-node fits, which slightly favour one-parameter templates; approximate analytic solutions of the damped stance (Saranlı et al., 2010), refined for damped and actuated legs (Seçer and Saranlı, 2018) and reviewed by Geyer and Saranlı (2018), could make exact fits cheap enough for every dataset. Datasets differ in quality: Baker et al. provide no COP, body mass or leg length, and Riglet et al. about 10 stances per condition. Trunk templates use a nominal trunk inertia and hip offset. The effective sample size in the BIC is a modelling choice, although the rankings were stable between 20 and 50 Hz and cross-validation does not depend on it.

## Acknowledgements

We thank the authors of the open datasets used here for making their data available. This work was partially supported by the Health Institutes of Türkiye (TÜSEB) under project number 40908.

## List of symbols and abbreviations

BIC: Bayesian information criterion
BW: body weight
COM, COP: centre of mass, centre of pressure
CPP, VPP: central pivot point, virtual pivot point
GRF: ground reaction force
SLIP: spring-loaded inverted pendulum
TD-SLIP: SLIP with a linearly decaying hip torque and leg damping
*c*, *c_bl_*: linear and bilinear leg damping constants (*c*^, *c*^*_bl_* dimensionless)
*k*, *k*^^^: leg stiffness, *k*^^^ = *kl*_0_*/mg*
*l*, *l*_0_: leg length, touchdown leg length
*t_s_*, *t_f_*: stance time, flight time
*τ*, *τ*^_0_: hip torque, dimensionless ramp amplitude *τ*_0_*/mgl*_0_
*x_ef_*: effective footprint

## Supplementary information

## Supplementary Materials and Methods

### Leg-axis laws

Spring *F_s_*: linear *k*(*l*_0_ *− l*); power law *kδ*_ref_(*δ/δ*_ref_)*^n^* with *δ* = *l*_0_ *− l*, *δ*_ref_ = 0.1*l*_0_; two-segment leg with a knee spring (segment length set for a 162^◦^ knee at touchdown, *k* the touchdown stiffness); stiffness switch at maximum compression, either with a force jump (*k_c_ → k_d_*) or force-continuous (*k_c_ → k_d_* and *l*_0_ *→ l*^′^ = *l_n_* + *k_c_*(*l*_0_ *−l_n_*)*/k_d_*). Damping *D*: none, linear *cl̇*, bilinear 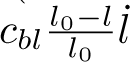, both, or compression-only *cl̇H*(*−l̇*).

### Hip laws

Ramp *τ*_0_(1 *− t/t_s_*); constant; half-stance constant; half-sine; force-proportional *ηF_r_l*(1 *− t/t_s_*) and *ηF_r_l*; hip torsional spring–damper *τ* = *k_h_*(*θ − θ*_0_) + *b_h_θ̇*; clock-torqued SLIP *τ* = *k_h_*(*θ − ϕ_c_*(*t*)) with *ϕ_c_*(*t*) = *θ_td_*(1 *−* 2*t/t_s_*) + *ϕ*_0_.

### Pivot laws

Point-mass virtual pivot at offsets (*x_vp_, r_vp_*) from the COM: *τ* = *F_r_l* (*x_vp_e_r,y_ − r_vp_e_r,x_*)*/*(*l* + *x_vp_e_r,x_* + *r_vp_e_r,y_*). Trunk templates (6-D state; hip 0.1*l*_0_ below the COM, trunk inertia *m*(0.25*l*_0_)^2^): the hip torque routes the GRF through a pivot *P* , *τ* = *−lF_r_*(***a*** *× **e**_r_*)*/*(***a*** *×* ***e****_t_*) with ***a*** = ***r****_f_ − P* ; *P* at the COM is the central pivot point, *P* on the trunk axis a trunk-fixed virtual pivot (touchdown pitch and pitch rate then solved with two extra periodicity constraints).

### Foot laws

Fixed pivot; pivot translating at a fitted speed or at the condition’s mean COP speed; foot mass *m_f_* = 0.06*m* joined to the body by the leg, with Hunt–Crossley normal contact *k_g_δ*^1.5^(1 + *αδ̇*), *α* = 1.5 s m^−1^, and a stick tangential contact; the measured GRF is then the contact force and the gait constraints apply to the whole-body COM.

### Information criteria

Per condition, *−*2 log *L* = *n* ln(SSE*/*2*N*) with *n* = 2*n*_eff_, *n*_eff_ = 2*f_c_t_s_*per component, *N* samples per component; BIC = *−*2 log *L* + *p* ln *n*; criteria are summed over conditions.

### Datasets

Fukuchi et al.: single-belt treadmill, forces 300 Hz, markers 150 Hz, pelvis markers for COM height. Carter et al.: instrumented treadmill, running on one plate at 1.5 kHz, 3-min trials; trials were streamed from the archive and processed on the fly. Baker et al.: GRF in body weights at about 120 Hz, one stride per file, no COP; flight time = half stride minus stance, nominal *l*_0_ = 1 m. Riglet et al.: two floor plates at 1 kHz, COM from the full-body model, both sessions pooled; the treadmill in this dataset is not instrumented. Code and processing scripts will be made publicly available before publication and can be shared with reviewers on request (see main text, Data and resource availability).

**Figure S1:**
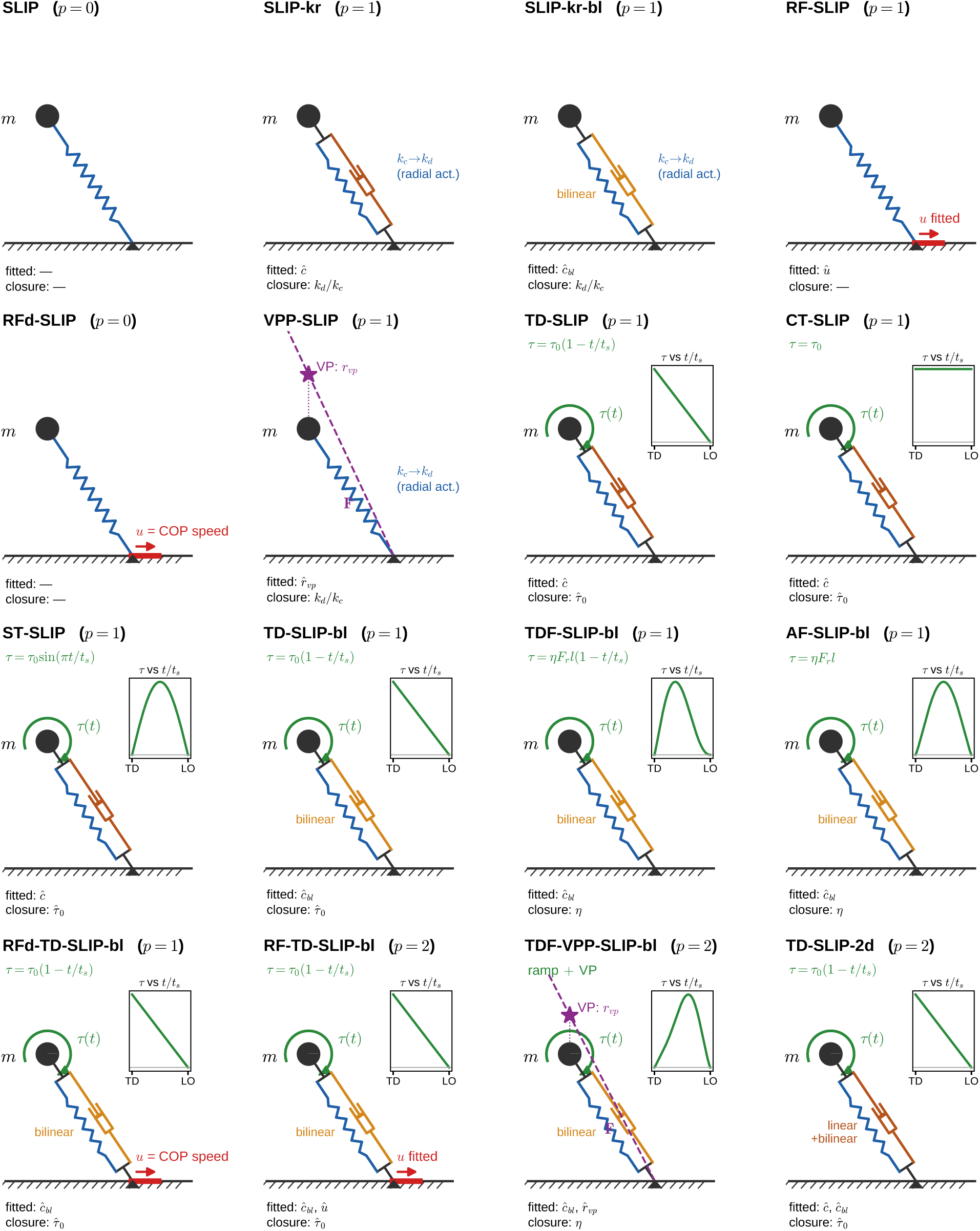
Templates (part 1). *p*, number of shape parameters fitted to the GRF; stiffness, touchdown state and one closure parameter are solved from the gait constraints.

**Figure S2:**
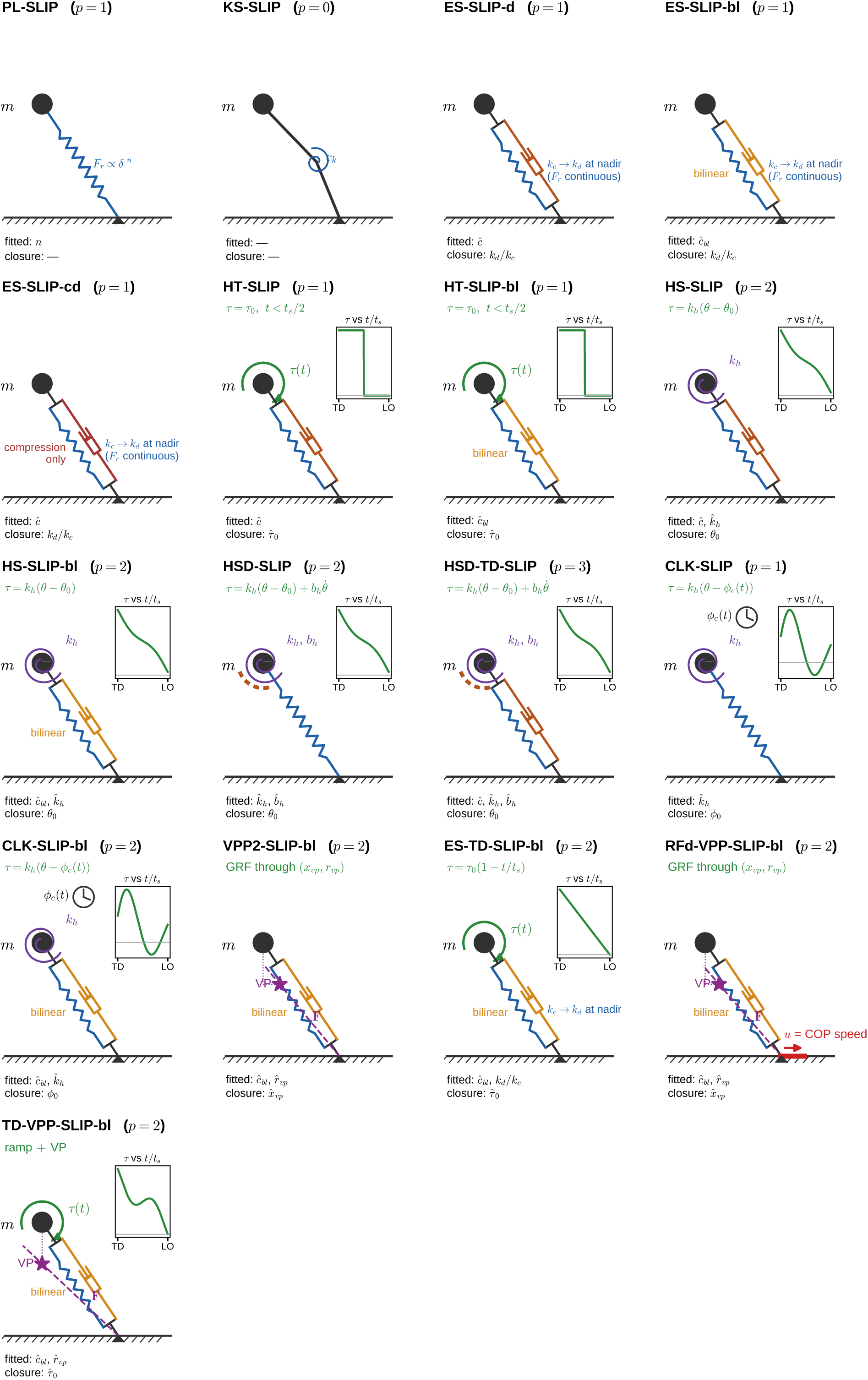
Templates (part 2).

**Figure S3:**
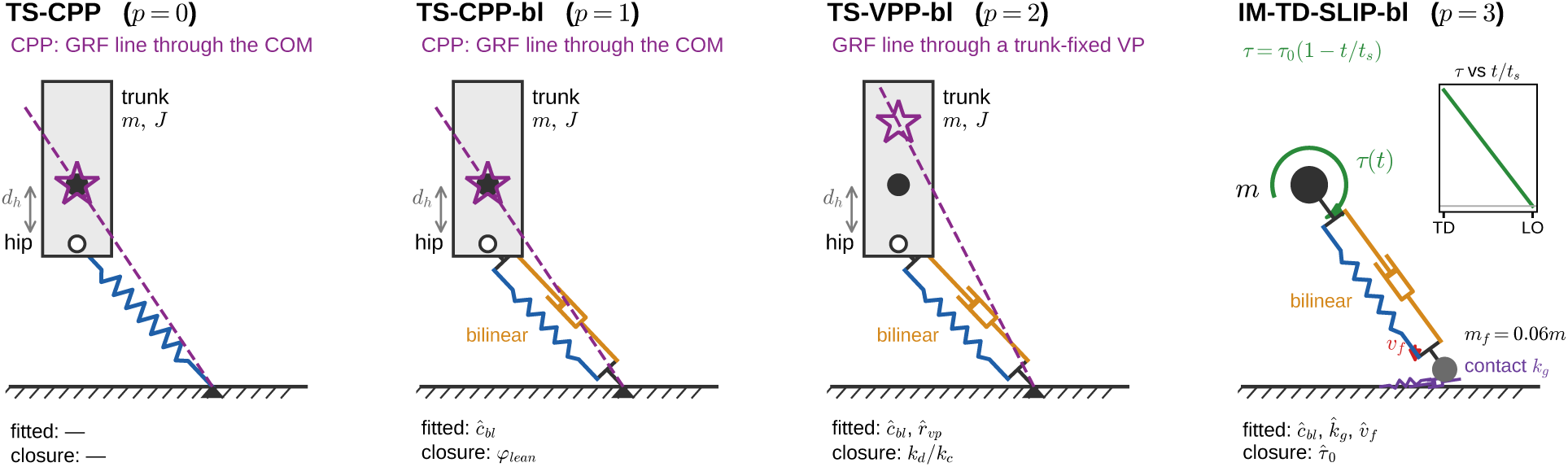
Templates (part 3): trunk models with the central pivot point and a trunk-fixed virtual pivot, and the foot-mass impact model.

---

**Table S1:** Datasets. Conditions entering the analysis (condition means; 414 human and 63 guinea-fowl conditions). Speeds in m s*^−^*^1^; COM, source of COM height at touchdown. Carter et al. runners were also recorded at 90 and 110% of habitual speed and Baker et al. runners at imposed cadences; these conditions are not included here. Goldsmith et al.: straight runs only; the exported COM height was shifted by one constant per bird to 1.06 hip heights at touchdown (Table S12 repeats the key fits with the exported height).

| Dataset | Setting | Subjects | Conditions | Steps | Speed | Force rate | COP | COM |
|---|---|---|---|---|---|---|---|---|
| Van Hooren et al. (Van Hooren and Meijer, 2024) | treadmill (split belt) | 19 | 95 | 28 512 | 2.78–5.00 | 1 kHz | yes | pelvis markers |
| Fukuchi et al. (Fukuchi et al., 2017) | treadmill | 39 | 101 | 8 482 | 2.50–4.50 | 300 Hz | yes | pelvis markers |
| Carter et al. (Carter et al., 2024) | treadmill | 50 | 96 | 64 131 | 2.08–4.25 | 1.5 kHz | yes | pelvis markers |
| Baker et al. (Baker et al., 2025) | treadmill | 15 | 50 | 2 695 | 2.00–4.00 | 120 Hz | no | nominal *l*<sub>0</sub> |
| Baker et al. (Baker et al., 2025) | overground | 15 | 15 | 1 175 | 2.23–3.52 | 120 Hz | no | nominal *l*<sub>0</sub> |
| Riglet et al. (Riglet et al., 2024) | overground | 30 | 57 | 647 | 2.08–4.78 | 1 kHz | yes | model COM |
| Daley & Birn-Jeffery (guinea fowl) (Daley and Birn-Jeffery, 2026) | overground | 13 | 37 | 435 | 1.44–3.98 | 500 Hz | foot | reconstructed COM |
| Goldsmith et al. (guinea fowl), high friction (Goldsmith et al., 2026) | overground | 5 | 13 | 225 | 1.36–2.47 | 800 Hz | yes | authors' COM, shifted |
| Goldsmith et al. (guinea fowl), low friction (Goldsmith et al., 2026) | overground | 6 | 13 | 247 | 1.33–2.26 | 800 Hz | yes | authors' COM, shifted |

**Table S2:**
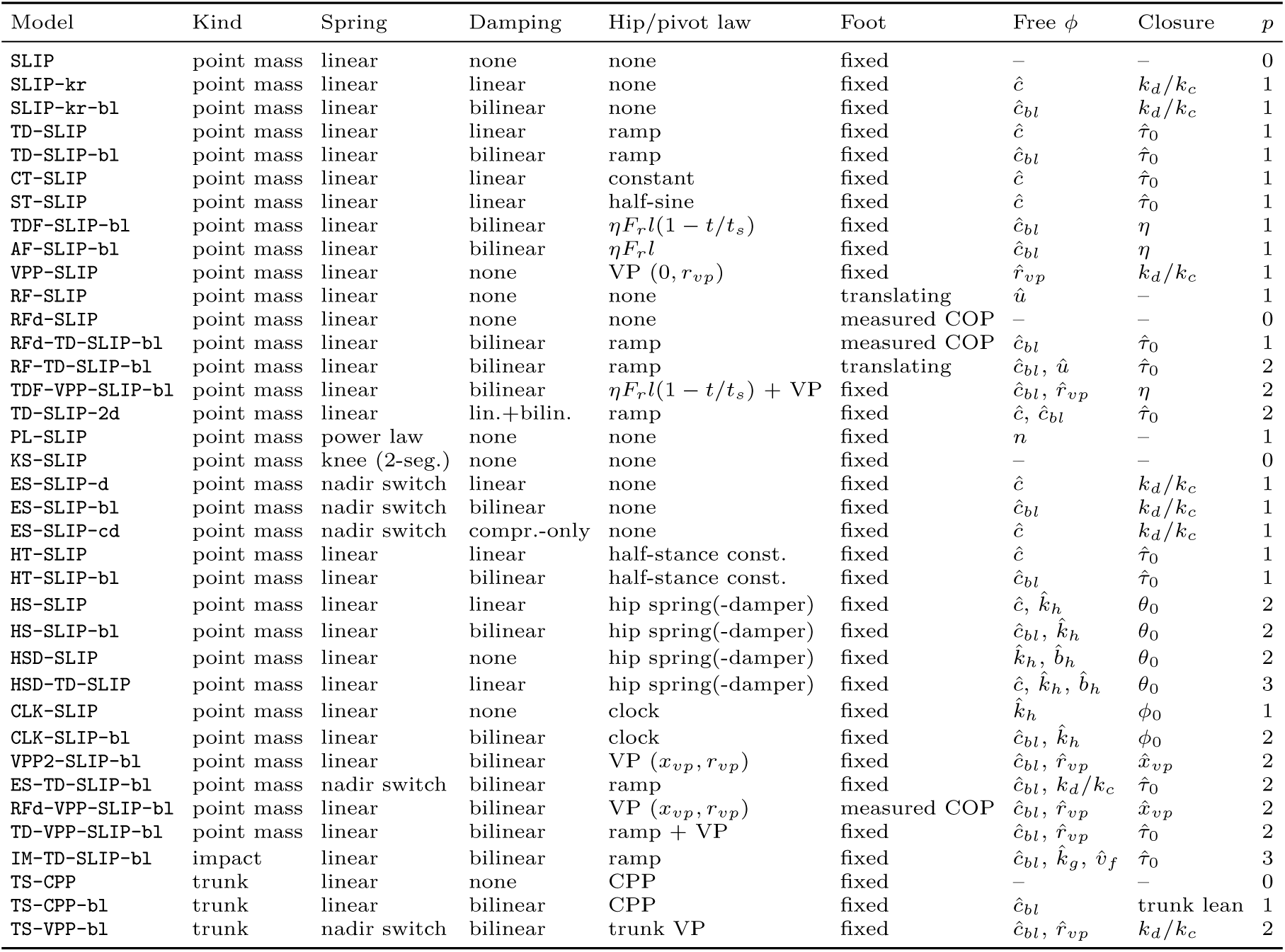
Template catalogue.

| Model | Kind | Spring | Damping | Hip/pivot law | Foot | Free *ϕ* | Closure | *p* |
|---|---|---|---|---|---|---|---|---|
| SLIP | point mass | linear | none | none | fixed | – | – | 0 |
| SLIP-kr | point mass | linear | linear | none | fixed | $\hat{c}$ | $k_d/k_c$ | 1 |
| SLIP-kr-bl | point mass | linear | bilinear | none | fixed | $\hat{c}_{bl}$ | $k_d/k_c$ | 1 |
| TD-SLIP | point mass | linear | linear | ramp | fixed | $\hat{c}$ | $\hat{\tau}_0$ | 1 |
| TD-SLIP-bl | point mass | linear | bilinear | ramp | fixed | $\hat{c}_{bl}$ | $\hat{\tau}_0$ | 1 |
| CT-SLIP | point mass | linear | linear | constant | fixed | $\hat{c}$ | $\hat{\tau}_0$ | 1 |
| ST-SLIP | point mass | linear | linear | half-sine | fixed | $\hat{c}$ | $\hat{\tau}_0$ | 1 |
| TDF-SLIP-bl | point mass | linear | bilinear | $\eta F_r l(1-t/t_s)$ | fixed | $\hat{c}_{bl}$ | *η* | 1 |
| AF-SLIP-bl | point mass | linear | bilinear | $\eta F_r l$ | fixed | $\hat{c}_{bl}$ | *η* | 1 |
| VPP-SLIP | point mass | linear | none | VP $(0, r_{vp})$ | fixed | $\hat{r}_{vp}$ | $k_d/k_c$ | 1 |
| RF-SLIP | point mass | linear | none | none | translating | $\hat{u}$ | – | 1 |
| RFd-SLIP | point mass | linear | none | none | measured COP | – | – | 0 |
| RFd-TD-SLIP-bl | point mass | linear | bilinear | ramp | measured COP | $\hat{c}_{bl}$ | $\hat{\tau}_0$ | 1 |
| RF-TD-SLIP-bl | point mass | linear | bilinear | ramp | translating | $\hat{c}_{bl}$, $\hat{u}$ | $\hat{\tau}_0$ | 2 |
| TDF-VPP-SLIP-bl | point mass | linear | bilinear | $\eta F_r l(1-t/t_s)$ + VP | fixed | $\hat{c}_{bl}$, $\hat{r}_{vp}$ | *η* | 2 |
| TD-SLIP-2d | point mass | linear | lin.+bilin. | ramp | fixed | $\hat{c}$, $\hat{c}_{bl}$ | $\hat{\tau}_0$ | 2 |
| PL-SLIP | point mass | power law | none | none | fixed | *n* | – | 1 |
| KS-SLIP | point mass | knee (2-seg.) | none | none | fixed | – | – | 0 |
| ES-SLIP-d | point mass | nadir switch | linear | none | fixed | $\hat{c}$ | $k_d/k_c$ | 1 |
| ES-SLIP-bl | point mass | nadir switch | bilinear | none | fixed | $\hat{c}_{bl}$ | $k_d/k_c$ | 1 |
| ES-SLIP-cd | point mass | nadir switch | compr.-only | none | fixed | $\hat{c}$ | $k_d/k_c$ | 1 |
| HT-SLIP | point mass | linear | linear | half-stance const. | fixed | $\hat{c}$ | $\hat{\tau}_0$ | 1 |
| HT-SLIP-bl | point mass | linear | bilinear | half-stance const. | fixed | $\hat{c}_{bl}$ | $\hat{\tau}_0$ | 1 |
| HS-SLIP | point mass | linear | linear | hip spring(-damper) | fixed | $\hat{c}$, $\hat{k}_h$ | $\theta_0$ | 2 |
| HS-SLIP-bl | point mass | linear | bilinear | hip spring(-damper) | fixed | $\hat{c}_{bl}$, $\hat{k}_h$ | $\theta_0$ | 2 |
| HSD-SLIP | point mass | linear | none | hip spring(-damper) | fixed | $\hat{k}_h$, $\hat{b}_h$ | $\theta_0$ | 2 |
| HSD-TD-SLIP | point mass | linear | linear | hip spring(-damper) | fixed | $\hat{c}$, $\hat{k}_h$, $\hat{b}_h$ | $\theta_0$ | 3 |
| CLK-SLIP | point mass | linear | none | clock | fixed | $\hat{k}_h$ | $\phi_0$ | 1 |
| CLK-SLIP-bl | point mass | linear | bilinear | clock | fixed | $\hat{c}_{bl}$, $\hat{k}_h$ | $\phi_0$ | 2 |
| VPP2-SLIP-bl | point mass | linear | bilinear | VP $(x_{vp}, r_{vp})$ | fixed | $\hat{c}_{bl}$, $\hat{r}_{vp}$ | $\hat{x}_{vp}$ | 2 |
| ES-TD-SLIP-bl | point mass | nadir switch | bilinear | ramp | fixed | $\hat{c}_{bl}$, $k_d/k_c$ | $\hat{\tau}_0$ | 2 |
| RFd-VPP-SLIP-bl | point mass | linear | bilinear | VP $(x_{vp}, r_{vp})$ | measured COP | $\hat{c}_{bl}$, $\hat{r}_{vp}$ | $\hat{x}_{vp}$ | 2 |
| TD-VPP-SLIP-bl | point mass | linear | bilinear | ramp + VP | fixed | $\hat{c}_{bl}$, $\hat{r}_{vp}$ | $\hat{\tau}_0$ | 2 |
| IM-TD-SLIP-bl | impact | linear | bilinear | ramp | fixed | $\hat{c}_{bl}$, $\hat{k}_g$, $\hat{v}_f$ | $\hat{\tau}_0$ | 3 |
| TS-CPP | trunk | linear | none | CPP | fixed | – | – | 0 |
| TS-CPP-bl | trunk | linear | bilinear | CPP | fixed | $\hat{c}_{bl}$ | trunk lean | 1 |
| TS-VPP-bl | trunk | nadir switch | bilinear | trunk VP | fixed | $\hat{c}_{bl}$, $\hat{r}_{vp}$ | $k_d/k_c$ | 2 |

**Table S3:** Free fits on Van Hooren et al. (95 conditions, exact refinement). RMSE in BW; ΔAIC and ΔBIC summed over conditions relative to the best model (*n*_eff_ from 50 Hz; second-to-last column 20 Hz); wins, conditions with the lowest BIC.

| Model | *p* | RMSE | RMSE<sub>*x*</sub> | RMSE<sub>*z*</sub> | ΔAIC | ΔBIC | ΔBIC<sub>20</sub> | wins |
|---|---|---|---|---|---|---|---|---|
| *Leg axis only* |  |  |  |  |  |  |  |  |
| ES-SLIP-d | 1 | 0.194 | 0.118 | 0.154 | 1668 | 1496 | 468 | 0 |
| ES-SLIP-bl | 1 | 0.203 | 0.106 | 0.173 | 1752 | 1580 | 502 | 0 |
| ES-SLIP-cd | 1 | 0.223 | 0.121 | 0.187 | 3084 | 2912 | 1035 | 0 |
| SLIP | 0 | 0.363 | 0.086 | 0.353 | 7049 | 6705 | 2422 | 0 |
| PL-SLIP | 1 | 0.353 | 0.095 | 0.340 | 6998 | 6826 | 2600 | 0 |
| KS-SLIP | 0 | 0.517 | 0.163 | 0.491 | 10370 | 10026 | 3750 | 0 |
| *Time-based hip torque* |  |  |  |  |  |  |  |  |
| TD-SLIP-2d | 2 | 0.160 | 0.080 | 0.138 | 0 | 0 | 0 | 51 |
| ES-TD-SLIP-bl | 2 | 0.169 | 0.078 | 0.150 | 471 | 471 | 188 | 13 |
| TD-SLIP-bl | 1 | 0.186 | 0.087 | 0.165 | 970 | 798 | 189 | 2 |
| HT-SLIP-bl | 1 | 0.189 | 0.091 | 0.166 | 1242 | 1070 | 298 | 1 |
| TDF-SLIP-bl | 1 | 0.195 | 0.095 | 0.171 | 1339 | 1167 | 337 | 3 |
| AF-SLIP-bl | 1 | 0.203 | 0.106 | 0.173 | 1665 | 1493 | 467 | 3 |
| TD-SLIP | 1 | 0.202 | 0.075 | 0.187 | 2084 | 1912 | 634 | 12 |
| HT-SLIP | 1 | 0.214 | 0.102 | 0.188 | 2612 | 2440 | 846 | 0 |
| CT-SLIP | 1 | 0.215 | 0.115 | 0.181 | 2643 | 2471 | 858 | 0 |
| ST-SLIP | 1 | 0.230 | 0.140 | 0.183 | 3259 | 3087 | 1105 | 0 |
| *State-based hip torque* |  |  |  |  |  |  |  |  |
| HS-SLIP-bl | 2 | 0.180 | 0.081 | 0.161 | 925 | 925 | 370 | 1 |
| CLK-SLIP-bl | 2 | 0.194 | 0.095 | 0.169 | 1570 | 1570 | 628 | 0 |
| HS-SLIP | 2 | 0.199 | 0.071 | 0.186 | 2152 | 2152 | 861 | 0 |
| HSD-TD-SLIP | 3 | 0.195 | 0.058 | 0.186 | 2186 | 2358 | 1073 | 7 |
| CLK-SLIP | 1 | 0.357 | 0.064 | 0.351 | 7073 | 6901 | 2630 | 0 |
| HSD-SLIP | 2 | 0.348 | 0.069 | 0.341 | 6974 | 6974 | 2790 | 0 |
| *Virtual / central pivot* |  |  |  |  |  |  |  |  |
| TD-VPP-SLIP-bl | 2 | 0.179 | 0.076 | 0.162 | 921 | 921 | 368 | 2 |
| TDF-VPP-SLIP-bl | 2 | 0.187 | 0.085 | 0.167 | 1260 | 1260 | 504 | 0 |
| VPP2-SLIP-bl | 2 | 0.188 | 0.087 | 0.167 | 1327 | 1327 | 531 | 0 |
| TS-VPP-bl | 2 | 0.191 | 0.086 | 0.170 | 1476 | 1476 | 590 | 0 |
| TS-CPP-bl | 1 | 0.208 | 0.111 | 0.176 | 2002 | 1830 | 602 | 0 |
| TS-CPP | 0 | 0.364 | 0.088 | 0.353 | 7062 | 6718 | 2427 | 0 |
| VPP-SLIP | 1 | 0.356 | 0.063 | 0.350 | 7034 | 6862 | 2615 | 0 |
| *Moving foot* |  |  |  |  |  |  |  |  |
| RF-TD-SLIP-bl | 2 | 0.181 | 0.075 | 0.165 | 982 | 982 | 393 | 0 |
| RFd-TD-SLIP-bl | 1 | 0.191 | 0.088 | 0.169 | 1369 | 1197 | 349 | 0 |
| RFd-VPP-SLIP-bl | 2 | 0.195 | 0.086 | 0.176 | 1662 | 1662 | 665 | 0 |
| RFd-SLIP | 0 | 0.361 | 0.068 | 0.355 | 6975 | 6631 | 2392 | 0 |
| RF-SLIP | 1 | 0.358 | 0.062 | 0.353 | 7100 | 6928 | 2641 | 0 |

**Table S4:** Cross-validated test RMSE on Van Hooren et al. (BW). Ratio, mean over held-out conditions of the MSE ratio to the best model.

|  |  | new speed |  | new runner (per speed) |  |  |  | one set for all |  |
|---|---|---|---|---|---|---|---|---|---|
| Model | *p* | RMSE | ratio | RMSE | RMSE<sub>*x*</sub> | RMSE<sub>*z*</sub> | ratio | RMSE | ratio |
| *Leg axis only* |  |  |  |  |  |  |  |  |  |
| ES-SLIP-d | 1 | 0.204 | 1.28 | 0.218 | 0.121 | 0.181 | 1.23 | 0.226 | 1.17 |
| ES-SLIP-bl | 1 | 0.226 | 1.52 | 0.235 | 0.109 | 0.209 | 1.40 | 0.242 | 1.31 |
| ES-SLIP-cd | 1 | 0.230 | 1.76 | 0.237 | 0.125 | 0.201 | 1.58 | 0.237 | 1.37 |
| PL-SLIP | 1 | 0.355 | 4.48 | 0.356 | 0.094 | 0.343 | 3.95 | 0.357 | 3.34 |
| SLIP | 0 | 0.363 | 4.65 | 0.363 | 0.086 | 0.353 | 4.08 | 0.363 | 3.46 |
| KS-SLIP | 0 | 0.517 | 9.93 | 0.517 | 0.163 | 0.491 | 8.73 | 0.517 | 7.51 |
| *Time-based hip torque* |  |  |  |  |  |  |  |  |  |
| TD-SLIP-2d | 2 | 0.187 | 1.00 | 0.202 | 0.078 | 0.187 | 1.00 | 0.216 | 1.00 |
| TD-SLIP | 1 | 0.212 | 1.46 | 0.215 | 0.075 | 0.202 | 1.28 | 0.219 | 1.13 |
| ES-TD-SLIP-bl | 2 | 0.413 | 3.32 | 0.218 | 0.082 | 0.202 | 1.17 | 0.242 | 1.23 |
| TD-SLIP-bl | 1 | 0.217 | 1.36 | 0.222 | 0.086 | 0.204 | 1.21 | 0.239 | 1.23 |
| HT-SLIP-bl | 1 | 0.218 | 1.38 | 0.222 | 0.088 | 0.203 | 1.23 | 0.240 | 1.25 |
| HT-SLIP | 1 | 0.225 | 1.66 | 0.226 | 0.099 | 0.203 | 1.43 | 0.230 | 1.28 |
| CT-SLIP | 1 | 0.225 | 1.62 | 0.229 | 0.119 | 0.196 | 1.43 | 0.232 | 1.26 |
| TDF-SLIP-bl | 1 | 0.224 | 1.46 | 0.230 | 0.096 | 0.209 | 1.32 | 0.248 | 1.34 |
| AF-SLIP-bl | 1 | 0.231 | 1.56 | 0.239 | 0.109 | 0.212 | 1.42 | 0.255 | 1.43 |
| ST-SLIP | 1 | 0.242 | 1.86 | 0.246 | 0.142 | 0.201 | 1.65 | 0.249 | 1.46 |
| *State-based hip torque* |  |  |  |  |  |  |  |  |  |
| HSD-TD-SLIP | 3 | 0.505 | 5.04 | 0.217 | 0.069 | 0.206 | 1.31 | 0.221 | 1.15 |
| HS-SLIP | 2 | 0.245 | 1.56 | 0.218 | 0.080 | 0.203 | 1.31 | 0.267 | 1.26 |
| HS-SLIP-bl | 2 | 0.213 | 1.32 | 0.220 | 0.086 | 0.203 | 1.21 | 0.242 | 1.27 |
| CLK-SLIP-bl | 2 | 0.224 | 1.47 | 0.230 | 0.098 | 0.208 | 1.33 | 0.244 | 1.33 |
| HSD-SLIP | 2 | 0.513 | 6.54 | 0.350 | 0.075 | 0.342 | 3.84 | 0.352 | 3.28 |
| CLK-SLIP | 1 | 0.357 | 4.53 | 0.357 | 0.064 | 0.351 | 3.97 | 0.357 | 3.36 |
| *Virtual / central pivot* |  |  |  |  |  |  |  |  |  |
| TD-VPP-SLIP-bl | 2 | 0.212 | 1.31 | 0.218 | 0.079 | 0.204 | 1.19 | 0.238 | 1.23 |
| TDF-VPP-SLIP-bl | 2 | 0.218 | 1.41 | 0.226 | 0.090 | 0.208 | 1.29 | 0.239 | 1.28 |
| VPP2-SLIP-bl | 2 | 0.220 | 1.43 | 0.227 | 0.093 | 0.207 | 1.30 | 0.240 | 1.29 |
| TS-VPP-bl | 2 | 0.220 | 1.43 | 0.234 | 0.094 | 0.214 | 1.37 | 0.235 | 1.23 |
| TS-CPP-bl | 1 | 0.309 | 2.36 | 0.258 | 0.116 | 0.232 | 1.72 | 0.281 | 1.72 |
| VPP-SLIP | 1 | 0.356 | 4.51 | 0.357 | 0.067 | 0.350 | 3.96 | 0.357 | 3.35 |
| TS-CPP | 0 | 0.364 | 4.66 | 0.364 | 0.088 | 0.353 | 4.08 | 0.364 | 3.47 |
| *Moving foot* |  |  |  |  |  |  |  |  |  |
| RF-TD-SLIP-bl | 2 | 0.213 | 1.32 | 0.219 | 0.079 | 0.204 | 1.19 | 0.230 | 1.16 |
| RFd-TD-SLIP-bl | 1 | 0.198 | 1.23 | 0.219 | 0.087 | 0.201 | 1.25 | 0.220 | 1.14 |
| RFd-VPP-SLIP-bl | 2 | 0.204 | 1.29 | 0.226 | 0.089 | 0.207 | 1.34 | 0.226 | 1.21 |
| RF-SLIP | 1 | 0.360 | 4.58 | 0.359 | 0.065 | 0.353 | 4.01 | 0.360 | 3.41 |
| RFd-SLIP | 0 | 0.361 | 4.62 | 0.361 | 0.068 | 0.355 | 4.05 | 0.361 | 3.44 |

**Table S5:**
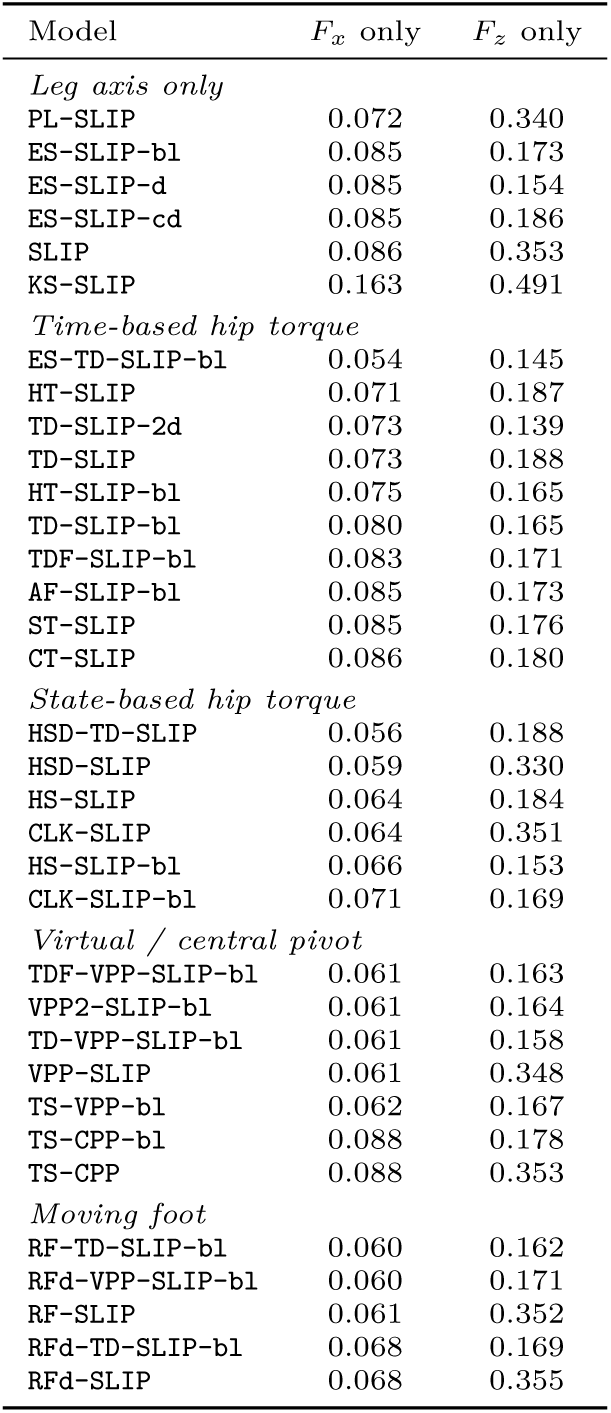
Best achievable component error when each force component is fitted alone (Van Hooren et al., RMSE in BW).

| Model | *F*<sub>*x*</sub> only | *F*<sub>*z*</sub> only |
|---|---|---|
| *Leg axis only* |  |  |
| PL-SLIP | 0.072 | 0.340 |
| ES-SLIP-bl | 0.085 | 0.173 |
| ES-SLIP-d | 0.085 | 0.154 |
| ES-SLIP-cd | 0.085 | 0.186 |
| SLIP | 0.086 | 0.353 |
| KS-SLIP | 0.163 | 0.491 |
| *Time-based hip torque* |  |  |
| ES-TD-SLIP-bl | 0.054 | 0.145 |
| HT-SLIP | 0.071 | 0.187 |
| TD-SLIP-2d | 0.073 | 0.139 |
| TD-SLIP | 0.073 | 0.188 |
| HT-SLIP-bl | 0.075 | 0.165 |
| TD-SLIP-bl | 0.080 | 0.165 |
| TDF-SLIP-bl | 0.083 | 0.171 |
| AF-SLIP-bl | 0.085 | 0.173 |
| ST-SLIP | 0.085 | 0.176 |
| CT-SLIP | 0.086 | 0.180 |
| *State-based hip torque* |  |  |
| HSD-TD-SLIP | 0.056 | 0.188 |
| HSD-SLIP | 0.059 | 0.330 |
| HS-SLIP | 0.064 | 0.184 |
| CLK-SLIP | 0.064 | 0.351 |
| HS-SLIP-bl | 0.066 | 0.153 |
| CLK-SLIP-bl | 0.071 | 0.169 |
| *Virtual / central pivot* |  |  |
| TDF-VPP-SLIP-bl | 0.061 | 0.163 |
| VPP2-SLIP-bl | 0.061 | 0.164 |
| TD-VPP-SLIP-bl | 0.061 | 0.158 |
| VPP-SLIP | 0.061 | 0.348 |
| TS-VPP-bl | 0.062 | 0.167 |
| TS-CPP-bl | 0.088 | 0.178 |
| TS-CPP | 0.088 | 0.353 |
| *Moving foot* |  |  |
| RF-TD-SLIP-bl | 0.060 | 0.162 |
| RFd-VPP-SLIP-bl | 0.060 | 0.171 |
| RF-SLIP | 0.061 | 0.352 |
| RFd-TD-SLIP-bl | 0.068 | 0.169 |
| RFd-SLIP | 0.068 | 0.355 |

**Table S6:** BIC excess per condition relative to the best model of each dataset (all templates). VH, Van Hooren; FU, Fukuchi; CA, Carter; BK, Baker treadmill; BKO, Baker overground; RG, Riglet; GF, Daley & Birn-Jeffery guinea fowl; GH and GL, Goldsmith et al. guinea fowl on high and low friction.

| Model | *p* | VH | FU | CA | BK | BKO | RG | GF | GH | GL |
|---|---|---|---|---|---|---|---|---|---|---|
| *Leg axis only* |  |  |  |  |  |  |  |  |  |  |
| ES-SLIP-bl | 1 | 16.6 | 14.4 | 9.7 | 10.3 | 7.9 | 32.2 | 30.7 | 47.5 | 41.2 |
| ES-SLIP-d | 1 | 15.7 | 12.9 | 9.0 | 32.4 | 30.1 | 13.9 | 26.1 | 45.9 | 37.8 |
| ES-SLIP-cd | 1 | 30.7 | 31.5 | 25.5 | 51.3 | 50.8 | 12.8 | 32.2 | 47.9 | 37.0 |
| PL-SLIP | 1 | 71.9 | 75.1 | 66.7 | 65.9 | 68.8 | 56.3 | 58.6 | 68.1 | 53.0 |
| SLIP | 0 | 70.6 | 72.8 | 64.1 | 69.5 | 71.9 | 60.3 | 58.7 | 70.3 | 57.5 |
| KS-SLIP | 0 | 105.5 | 135.1 | 110.5 | 138.3 | 150.0 | 101.0 | 65.2 | 110.4 | 106.7 |
| *Time-based hip torque* |  |  |  |  |  |  |  |  |  |  |
| TD-SLIP-2d | 2 | **0** | **0** | **0** | 9.3 | 7.1 | 1.5 | 30.4 | 47.1 | 37.2 |
| HT-SLIP-bl | 1 | 11.3 | 8.0 | 6.5 | 5.9 | 3.4 | 26.8 | 22.4 | 37.1 | 31.7 |
| TD-SLIP-bl | 1 | 8.4 | 6.6 | 4.8 | 5.3 | 3.1 | 26.9 | 31.7 | 47.4 | 39.6 |
| TDF-SLIP-bl | 1 | 12.3 | 9.7 | 6.0 | 4.0 | 1.0 | 30.1 | 28.4 | 44.7 | 38.4 |
| ES-TD-SLIP-bl | 2 | 5.0 | 7.2 | 5.7 | 7.9 | 6.1 | 23.7 | 33.7 | 50.0 | 42.8 |
| AF-SLIP-bl | 1 | 15.7 | 13.4 | 8.5 | 6.4 | 4.0 | 32.1 | 37.4 | 53.8 | 45.2 |
| HT-SLIP | 1 | 25.7 | 27.5 | 21.6 | 45.6 | 44.9 | 6.6 | 23.6 | 38.8 | 25.4 |
| TD-SLIP | 1 | 20.1 | 22.1 | 16.8 | 45.0 | 44.4 | **0** | 34.0 | 50.7 | 36.9 |
| CT-SLIP | 1 | 26.0 | 25.4 | 20.6 | 46.4 | 46.4 | 8.7 | 48.7 | 63.0 | 49.9 |
| ST-SLIP | 1 | 32.5 | 30.7 | 23.9 | 47.6 | 47.2 | 15.7 | 48.7 | 63.0 | 49.4 |
| *State-based hip torque* |  |  |  |  |  |  |  |  |  |  |
| HS-SLIP-bl | 2 | 9.7 | 9.1 | 7.9 | 9.4 | 7.0 | 28.9 | 18.9 | 23.5 | 18.4 |
| CLK-SLIP-bl | 2 | 16.5 | 14.2 | 12.0 | 10.7 | 9.6 | 34.2 | 30.2 | 37.2 | 32.6 |
| HS-SLIP | 2 | 22.7 | 24.7 | 20.3 | 49.2 | 48.3 | 3.6 | 26.8 | 33.0 | 23.3 |
| HSD-TD-SLIP | 3 | 24.8 | 30.5 | 25.1 | 55.7 | 54.6 | 8.1 | 31.6 | 38.2 | 26.4 |
| CLK-SLIP | 1 | 72.6 | 75.2 | 66.9 | 72.4 | 74.7 | 62.8 | 51.7 | 51.1 | 40.6 |
| HSD-SLIP | 2 | 73.4 | 77.6 | 69.4 | 76.1 | 78.3 | 63.8 | 55.9 | 55.3 | 45.9 |
| *Virtual / central pivot* |  |  |  |  |  |  |  |  |  |  |
| TD-VPP-SLIP-bl | 2 | 9.7 | 7.0 | 7.5 | 7.2 | 4.4 | 29.2 | 1.7 | 1.7 | 1.4 |
| TDF-VPP-SLIP-bl | 2 | 13.3 | 10.4 | 8.6 | 6.4 | 3.3 | 32.1 | 0.8 | **0** | **0** |
| VPP2-SLIP-bl | 2 | 14.0 | 11.1 | 9.0 | 6.6 | 3.5 | 32.3 | 0.8 | 0.1 | 0.1 |
| TS-VPP-bl | 2 | 15.5 | 13.3 | 10.7 | 11.0 | 8.6 | 32.8 | 3.7 | 12.2 | 6.4 |
| TS-CPP-bl | 1 | 19.3 | 16.9 | 10.7 | 14.7 | 11.5 | 32.7 | 50.2 | 63.8 | 50.1 |
| VPP-SLIP | 1 | 72.2 | 74.9 | 66.7 | 72.2 | 74.4 | 62.4 | 50.4 | 47.8 | 38.1 |
| TS-CPP | 0 | 70.7 | 72.9 | 64.1 | 69.8 | 72.3 | 60.3 | 60.2 | 76.2 | 59.4 |
| *Moving foot* |  |  |  |  |  |  |  |  |  |  |
| RF-TD-SLIP-bl | 2 | 10.3 | 6.2 | 7.8 | **0** | **0** | 30.2 | 7.9 | 15.1 | 14.8 |
| RFd-VPP-SLIP-bl | 2 | 17.5 | 11.6 | 10.7 | 6.6 | 3.5 | 36.3 | **0** | 4.4 | 4.0 |
| RFd-TD-SLIP-bl | 1 | 12.6 | 5.7 | 7.2 | 5.3 | 3.1 | 30.5 | 21.3 | 29.3 | 28.8 |
| RFd-SLIP | 0 | 69.8 | 71.6 | 63.5 | 69.5 | 71.9 | 60.8 | 54.7 | 58.1 | 50.3 |
| RF-SLIP | 1 | 72.9 | 75.1 | 67.0 | 70.0 | 72.2 | 63.5 | 53.1 | 56.9 | 46.7 |

**Table S7:** Cross-validated median test RMSE (BW): one parameter set fitted on all other runners of the dataset (bold, best).

| Model | *p* | VH | FU | CA | BK | BKO | RG | GF | GH | GL |
|---|---|---|---|---|---|---|---|---|---|---|
| *Leg axis only* |  |  |  |  |  |  |  |  |  |  |
| ES-SLIP-bl | 1 | 0.218 | 0.203 | 0.215 | 0.216 | 0.199 | 0.248 | 0.241 | 0.248 | 0.190 |
| ES-SLIP-d | 1 | 0.213 | 0.189 | 0.209 | 0.250 | 0.232 | 0.202 | 0.221 | 0.223 | 0.182 |
| ES-SLIP-cd | 1 | 0.233 | 0.222 | 0.245 | 0.301 | 0.275 | 0.203 | 0.263 | 0.228 | 0.204 |
| PL-SLIP | 1 | 0.358 | 0.341 | 0.366 | 0.309 | 0.305 | 0.301 | 0.431 | 0.328 | 0.271 |
| SLIP | 0 | 0.368 | 0.343 | 0.367 | 0.334 | 0.322 | 0.329 | 0.457 | 0.357 | 0.285 |
| KS-SLIP | 0 | 0.517 | 0.518 | 0.546 | 0.580 | 0.579 | 0.415 | 0.535 | 0.451 | 0.416 |
| *Time-based hip torque* |  |  |  |  |  |  |  |  |  |  |
| TD-SLIP-2d | 2 | **0.199** | 0.207 | **0.201** | 0.213 | 0.191 | **0.174** | 0.277 | 0.276 | 0.194 |
| HT-SLIP-bl | 1 | 0.217 | 0.190 | 0.214 | 0.206 | 0.186 | 0.238 | 0.246 | 0.280 | 0.160 |
| TD-SLIP-bl | 1 | 0.216 | 0.188 | 0.211 | 0.207 | 0.192 | 0.242 | 0.286 | 0.290 | 0.186 |
| TDF-SLIP-bl | 1 | 0.222 | 0.196 | 0.214 | 0.206 | 0.195 | 0.243 | 0.254 | 0.278 | 0.183 |
| ES-TD-SLIP-bl | 2 | 0.216 | 0.198 | 0.210 | 0.209 | 0.192 | 0.241 | 0.249 | 0.234 | 0.184 |
| AF-SLIP-bl | 1 | 0.226 | 0.200 | 0.218 | 0.214 | 0.199 | 0.247 | 0.308 | 0.297 | 0.206 |
| HT-SLIP | 1 | 0.223 | 0.219 | 0.236 | 0.291 | 0.260 | 0.182 | 0.231 | 0.263 | 0.152 |
| TD-SLIP | 1 | 0.212 | 0.205 | 0.228 | 0.285 | 0.257 | 0.185 | 0.275 | 0.285 | 0.187 |
| CT-SLIP | 1 | 0.222 | 0.212 | 0.234 | 0.291 | 0.265 | 0.195 | 0.360 | 0.330 | 0.228 |
| ST-SLIP | 1 | 0.236 | 0.224 | 0.242 | 0.292 | 0.271 | 0.209 | 0.348 | 0.331 | 0.227 |
| *State-based hip torque* |  |  |  |  |  |  |  |  |  |  |
| HS-SLIP-bl | 2 | 0.217 | 0.200 | 0.209 | 0.213 | 0.189 | 0.240 | 0.198 | 0.214 | 0.149 |
| CLK-SLIP-bl | 2 | 0.220 | 0.195 | 0.214 | 0.219 | 0.191 | 0.241 | 0.235 | 0.264 | 0.170 |
| HS-SLIP | 2 | 0.216 | 0.207 | 0.226 | 0.285 | 0.257 | 0.177 | 0.237 | 0.234 | 0.151 |
| HSD-TD-SLIP | 3 | 0.215 | 0.207 | 0.228 | 0.289 | 0.262 | 0.179 | 0.257 | 0.235 | 0.150 |
| CLK-SLIP | 1 | 0.356 | 0.343 | 0.362 | 0.331 | 0.319 | 0.327 | 0.376 | 0.315 | 0.216 |
| HSD-SLIP | 2 | 0.342 | 0.342 | 0.363 | 0.331 | 0.319 | 0.325 | 0.384 | 0.291 | 0.224 |
| *Virtual / central pivot* |  |  |  |  |  |  |  |  |  |  |
| TD-VPP-SLIP-bl | 2 | 0.215 | **0.186** | 0.208 | 0.212 | 0.185 | 0.238 | 0.143 | 0.170 | 0.132 |
| TDF-VPP-SLIP-bl | 2 | 0.217 | 0.193 | 0.211 | 0.209 | 0.185 | 0.243 | 0.146 | 0.170 | 0.131 |
| VPP2-SLIP-bl | 2 | 0.217 | 0.194 | 0.212 | 0.210 | **0.184** | 0.244 | **0.142** | 0.170 | 0.135 |
| TS-VPP-bl | 2 | 0.212 | 0.192 | 0.212 | 0.216 | 0.188 | 0.248 | 0.177 | 0.178 | **0.103** |
| TS-CPP-bl | 1 | 0.239 | 0.227 | 0.227 | 0.235 | 0.197 | 0.250 | 0.363 | 0.284 | 0.232 |
| VPP-SLIP | 1 | 0.355 | 0.343 | 0.363 | 0.330 | 0.319 | 0.327 | 0.364 | 0.267 | 0.211 |
| TS-CPP | 0 | 0.368 | 0.343 | 0.367 | 0.336 | 0.323 | 0.328 | 0.466 | 0.357 | 0.292 |
| *Moving foot* |  |  |  |  |  |  |  |  |  |  |
| RF-TD-SLIP-bl | 2 | 0.206 | 0.187 | 0.207 | **0.196** | 0.185 | 0.240 | 0.187 | 0.161 | 0.120 |
| RFd-VPP-SLIP-bl | 2 | 0.206 | 0.194 | 0.215 | 0.210 | **0.184** | 0.253 | 0.144 | **0.093** | 0.112 |
| RFd-TD-SLIP-bl | 1 | 0.202 | 0.188 | 0.213 | 0.207 | 0.192 | 0.248 | 0.228 | 0.184 | 0.166 |
| RFd-SLIP | 0 | 0.359 | 0.345 | 0.364 | 0.334 | 0.322 | 0.334 | 0.425 | 0.314 | 0.262 |
| RF-SLIP | 1 | 0.360 | 0.344 | 0.363 | 0.323 | 0.310 | 0.329 | 0.388 | 0.292 | 0.230 |

**Table S8:** 4 m s*^−^*^1^ subset (19 runners) including the foot-mass impact model: RMSE in BW over the whole stance and separately for the first 15% (impact) and the rest; ΔBIC summed.

|  |  | whole stance |  |  |  | first 15% |  | rest |  |
|---|---|---|---|---|---|---|---|---|---|
| Model | *p* | RMSE | *F<sub>x</sub>* | *F<sub>z</sub>* | ΔBIC | *F<sub>x</sub>* | *F<sub>z</sub>* | *F<sub>x</sub>* | *F<sub>z</sub>* |
| TD-SLIP-2d | 2 | 0.170 | 0.084 | 0.147 | 0 | 0.139 | 0.270 | 0.070 | 0.112 |
| ES-TD-SLIP-bl | 2 | 0.179 | 0.082 | 0.159 | 72 | 0.172 | 0.284 | 0.051 | 0.124 |
| IM-TD-SLIP-bl | 3 | 0.169 | 0.079 | 0.149 | 75 | 0.163 | 0.231 | 0.052 | 0.129 |
| TD-SLIP-bl | 1 | 0.199 | 0.091 | 0.177 | 145 | 0.144 | 0.308 | 0.078 | 0.141 |
| ES-SLIP-d | 1 | 0.206 | 0.125 | 0.163 | 256 | 0.217 | 0.294 | 0.100 | 0.127 |
| TD-SLIP | 1 | 0.210 | 0.079 | 0.194 | 305 | 0.130 | 0.355 | 0.066 | 0.148 |
| HSD-TD-SLIP | 3 | 0.203 | 0.062 | 0.193 | 391 | 0.097 | 0.358 | 0.053 | 0.145 |
| SLIP | 0 | 0.363 | 0.089 | 0.352 | 1094 | 0.143 | 0.534 | 0.075 | 0.308 |
| SLIP-kr-bl | 1 | 0.363 | 0.089 | 0.352 | 1164 | 0.143 | 0.533 | 0.075 | 0.308 |
| SLIP-kr | 1 | 0.363 | 0.089 | 0.352 | 1165 | 0.143 | 0.534 | 0.075 | 0.308 |

**Table S9:**
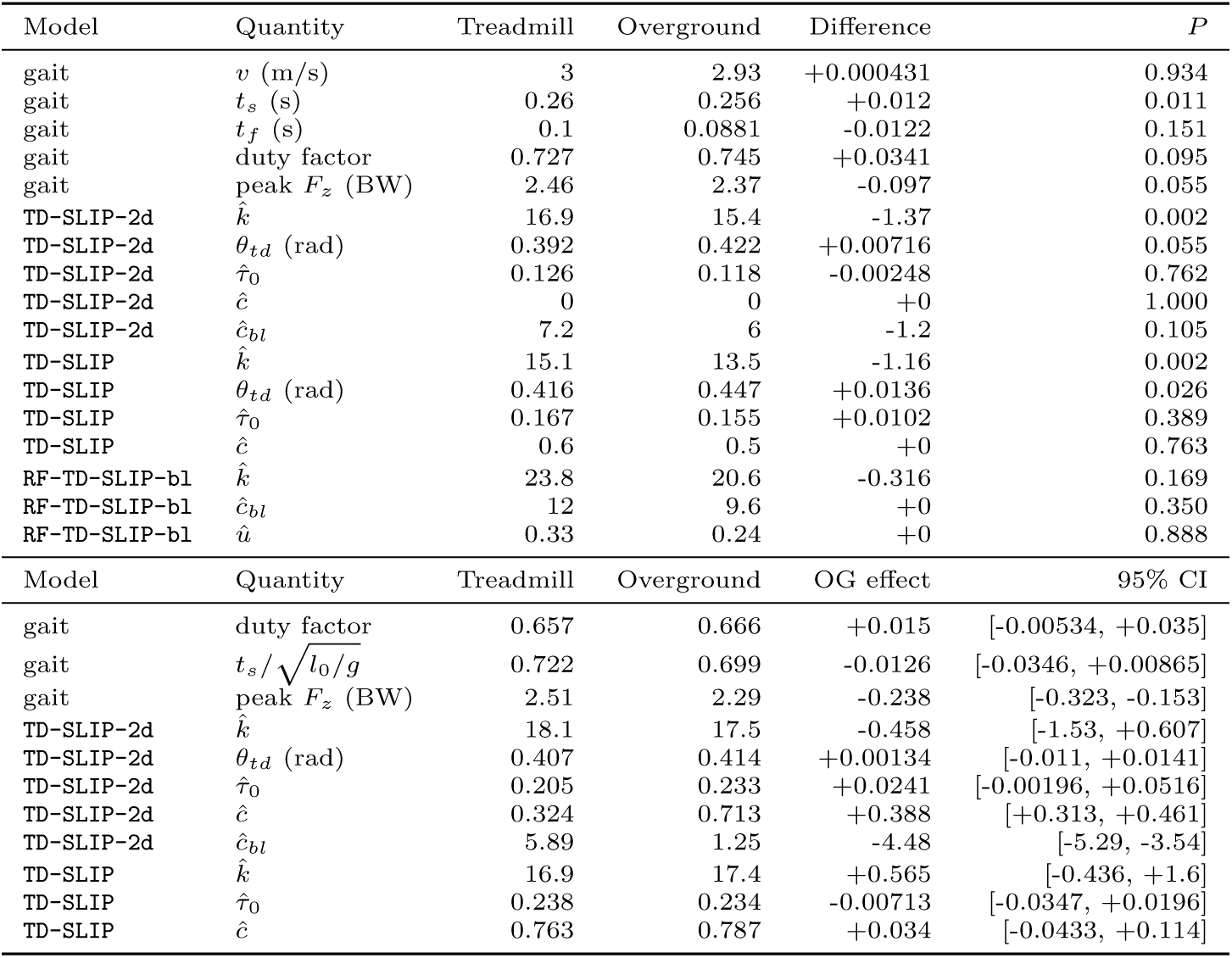
Treadmill versus overground. Top: same runners (Baker et al., *n* = 15 pairs; medians, median paired difference, Wilcoxon *P*). Bottom: Riglet et al. (overground) vs Van Hooren, Fukuchi and Carter (treadmill), speed-matched; overground effect adjusted for dimensionless speed with 95% runner-clustered bootstrap interval.

| Model | Quantity | Treadmill | Overground | Difference | *P* |
|---|---|---|---|---|---|
| gait | *v* (m/s) | 3 | 2.93 | +0.000431 | 0.934 |
| gait | *t*<sub>*s*</sub> (s) | 0.26 | 0.256 | +0.012 | 0.011 |
| gait | *t*<sub>*f*</sub> (s) | 0.1 | 0.0881 | -0.0122 | 0.151 |
| gait | duty factor | 0.727 | 0.745 | +0.0341 | 0.095 |
| gait | peak *F*<sub>*z*</sub> (BW) | 2.46 | 2.37 | -0.097 | 0.055 |
| TD-SLIP-2d | *k̂* | 16.9 | 15.4 | -1.37 | 0.002 |
| TD-SLIP-2d | *θ*<sub>*td*</sub> (rad) | 0.392 | 0.422 | +0.00716 | 0.055 |
| TD-SLIP-2d | *τ̂*<sub>0</sub> | 0.126 | 0.118 | -0.00248 | 0.762 |
| TD-SLIP-2d | *ĉ* | 0 | 0 | +0 | 1.000 |
| TD-SLIP-2d | *ĉ*<sub>*bl*</sub> | 7.2 | 6 | -1.2 | 0.105 |
| TD-SLIP | *k̂* | 15.1 | 13.5 | -1.16 | 0.002 |
| TD-SLIP | *θ*<sub>*td*</sub> (rad) | 0.416 | 0.447 | +0.0136 | 0.026 |
| TD-SLIP | *τ̂*<sub>0</sub> | 0.167 | 0.155 | +0.0102 | 0.389 |
| TD-SLIP | *ĉ* | 0.6 | 0.5 | +0 | 0.763 |
| RF-TD-SLIP-bl | *k̂* | 23.8 | 20.6 | -0.316 | 0.169 |
| RF-TD-SLIP-bl | *ĉ*<sub>*bl*</sub> | 12 | 9.6 | +0 | 0.350 |
| RF-TD-SLIP-bl | *û* | 0.33 | 0.24 | +0 | 0.888 |
| Model | Quantity | Treadmill | Overground | OG effect | 95% CI |
| gait | duty factor | 0.657 | 0.666 | +0.015 | [-0.00534, +0.035] |
| gait | $t_s/\sqrt{l_0/g}$ | 0.722 | 0.699 | -0.0126 | [-0.0346, +0.00865] |
| gait | peak *F*<sub>*z*</sub> (BW) | 2.51 | 2.29 | -0.238 | [-0.323, -0.153] |
| TD-SLIP-2d | *k̂* | 18.1 | 17.5 | -0.458 | [-1.53, +0.607] |
| TD-SLIP-2d | *θ*<sub>*td*</sub> (rad) | 0.407 | 0.414 | +0.00134 | [-0.011, +0.0141] |
| TD-SLIP-2d | *τ̂*<sub>0</sub> | 0.205 | 0.233 | +0.0241 | [-0.00196, +0.0516] |
| TD-SLIP-2d | *ĉ* | 0.324 | 0.713 | +0.388 | [+0.313, +0.461] |
| TD-SLIP-2d | *ĉ*<sub>*bl*</sub> | 5.89 | 1.25 | -4.48 | [-5.29, -3.54] |
| TD-SLIP | *k̂* | 16.9 | 17.4 | +0.565 | [-0.436, +1.6] |
| TD-SLIP | *τ̂*<sub>0</sub> | 0.238 | 0.234 | -0.00713 | [-0.0347, +0.0196] |
| TD-SLIP | *ĉ* | 0.763 | 0.787 | +0.034 | [-0.0433, +0.114] |

**Table S10:** Reproduction of the 2012 errors on the digitized curves (MSE in N^2^, *m* = 64.5 kg, *l*_0_ = 1 m); stride-normalised MSE multiplies the stance MSE by *t_s_/*(*t_s_* + *t_f_*), as in the 2012 study.

|  | 2012 abstract |  | present, stride-norm. |  | difference |  | present, stance MSE |  |
|---|---|---|---|---|---|---|---|---|
| Data | SLIP | TD-SLIP | SLIP | TD-SLIP | SLIP | TD-SLIP | SLIP | TD-SLIP |
| Kram 1998, 3 m/s (overground) | 44657 | 12424 | 44458 | 12173 | -0.4% | -2.0% | 66738 | 18273 |
| Hamill 1983, 4 m/s | 48584 | 22413 | 48116 | 22211 | -1.0% | -0.9% | 79752 | 36815 |
| Hamill 1983, 5 m/s | 68065 | 29960 | 67448 | 29877 | -0.9% | -0.3% | 118533 | 52507 |
| Hamill 1983, 6 m/s | 87664 | 32837 | 87068 | 32631 | -0.7% | -0.6% | 166780 | 62505 |

**Table S11:**
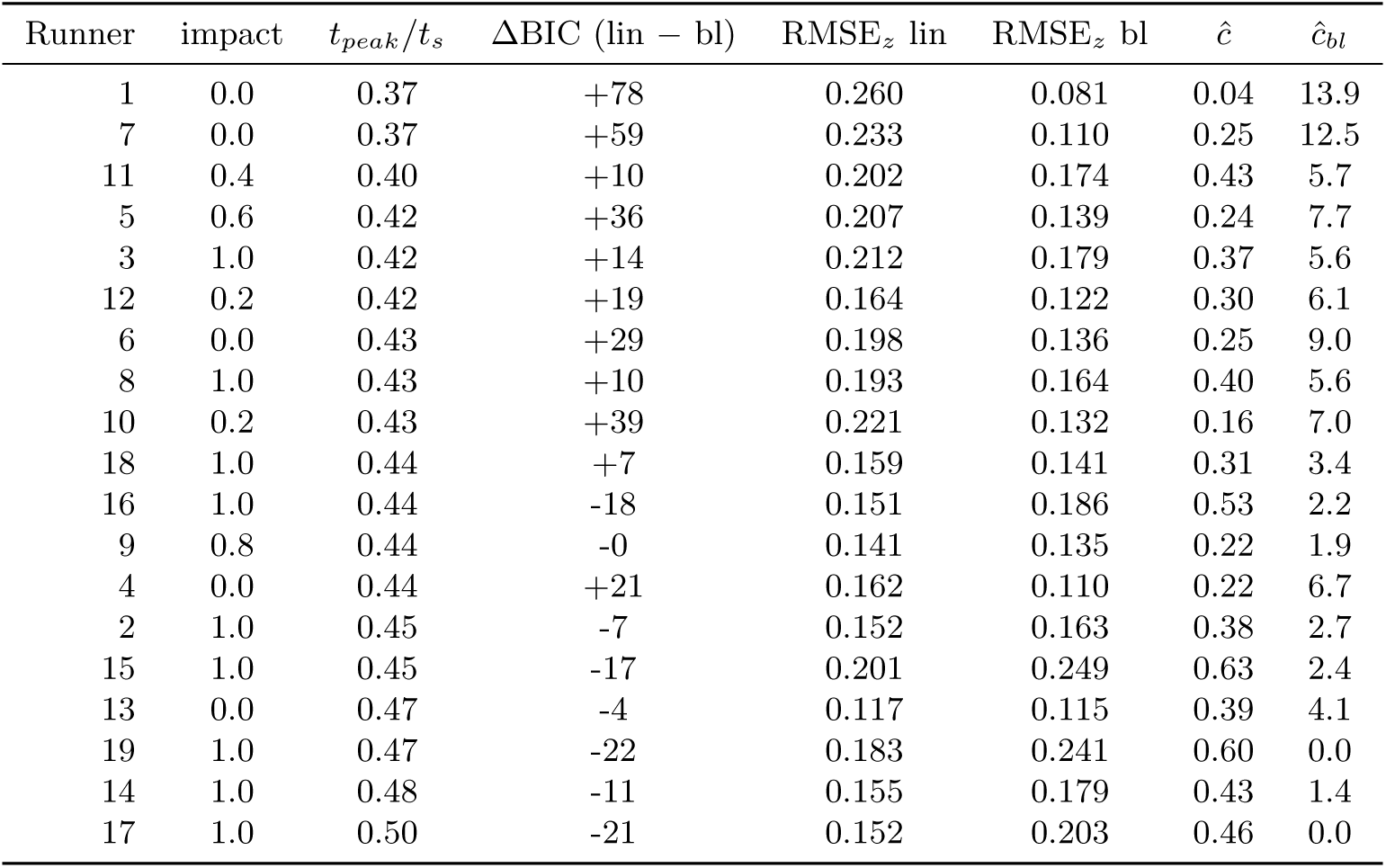
Per-runner relation between vertical-force shape and damping (Van Hooren et al., means over speeds). Impact, fraction of speeds with a local maximum before the main peak; ΔBIC *>* 0 favours bilinear over linear TD-SLIP; last columns, median damping constants of TD-SLIP-2d.

| Runner | impact | t<sub>peak</sub>/t<sub>s</sub> | ΔBIC (lin − bl) | RMSE<sub>z</sub> lin | RMSE<sub>z</sub> bl | ĉ | ĉ<sub>bl</sub> |
|---|---|---|---|---|---|---|---|
| 1 | 0.0 | 0.37 | +78 | 0.260 | 0.081 | 0.04 | 13.9 |
| 7 | 0.0 | 0.37 | +59 | 0.233 | 0.110 | 0.25 | 12.5 |
| 11 | 0.4 | 0.40 | +10 | 0.202 | 0.174 | 0.43 | 5.7 |
| 5 | 0.6 | 0.42 | +36 | 0.207 | 0.139 | 0.24 | 7.7 |
| 3 | 1.0 | 0.42 | +14 | 0.212 | 0.179 | 0.37 | 5.6 |
| 12 | 0.2 | 0.42 | +19 | 0.164 | 0.122 | 0.30 | 6.1 |
| 6 | 0.0 | 0.43 | +29 | 0.198 | 0.136 | 0.25 | 9.0 |
| 8 | 1.0 | 0.43 | +10 | 0.193 | 0.164 | 0.40 | 5.6 |
| 10 | 0.2 | 0.43 | +39 | 0.221 | 0.132 | 0.16 | 7.0 |
| 18 | 1.0 | 0.44 | +7 | 0.159 | 0.141 | 0.31 | 3.4 |
| 16 | 1.0 | 0.44 | -18 | 0.151 | 0.186 | 0.53 | 2.2 |
| 9 | 0.8 | 0.44 | -0 | 0.141 | 0.135 | 0.22 | 1.9 |
| 4 | 0.0 | 0.44 | +21 | 0.162 | 0.110 | 0.22 | 6.7 |
| 2 | 1.0 | 0.45 | -7 | 0.152 | 0.163 | 0.38 | 2.7 |
| 15 | 1.0 | 0.45 | -17 | 0.201 | 0.249 | 0.63 | 2.4 |
| 13 | 0.0 | 0.47 | -4 | 0.117 | 0.115 | 0.39 | 4.1 |
| 19 | 1.0 | 0.47 | -22 | 0.183 | 0.241 | 0.60 | 0.0 |
| 14 | 1.0 | 0.48 | -11 | 0.155 | 0.179 | 0.43 | 1.4 |
| 17 | 1.0 | 0.50 | -21 | 0.152 | 0.203 | 0.46 | 0.0 |

**Table S12:** Second guinea-fowl dataset (Goldsmith et al.): sensitivity to the COM height. For each surface, BIC excess per condition among the nine templates listed (0, best), fore-aft RMSE (BW) and median fitted pivot height above the COM (*l*_0_), with the COM height shifted to 1.06 hip heights at touchdown (used in the main text) and as exported (about 0.08 m higher).

|  | shifted COM |  |  |  |  |  | exported COM |  |  |  |  |  |
|---|---|---|---|---|---|---|---|---|---|---|---|---|
|  | high friction |  |  | low friction |  |  | high friction |  |  | low friction |  |  |
| Template | ΔBIC | *x* | pivot | ΔBIC | *x* | pivot | ΔBIC | *x* | pivot | ΔBIC | *x* | pivot |
| TDF-VPP-SLIP-bl | **0** | 0.043 | 1.15 | **0** | 0.037 | 0.85 | 0.2 | 0.038 | 0.55 | 0.1 | 0.036 | 0.40 |
| VPP2-SLIP-bl | 0.1 | 0.042 | 1.30 | 0.1 | 0.037 | 1.00 | **0** | 0.038 | 0.70 | **0** | 0.034 | 0.40 |
| TD-VPP-SLIP-bl | 1.7 | 0.049 | 1.15 | 1.4 | 0.043 | 1.00 | 3.5 | 0.049 | 0.55 | 2.0 | 0.044 | 0.40 |
| RFd-VPP-SLIP-bl | 4.4 | 0.042 | 0.85 | 4.0 | 0.038 | 0.85 | 7.2 | 0.039 | 0.25 | 5.5 | 0.033 | 0.25 |
| TS-VPP-bl | 12.2 | 0.074 | 0.55 | 6.4 | 0.032 | 0.55 | 1.5 | 0.033 | 0.40 | 0.8 | 0.029 | 0.40 |
| RF-TD-SLIP-bl | 15.1 | 0.070 | – | 14.8 | 0.058 | – | 11.0 | 0.065 | – | 8.2 | 0.052 | – |
| TD-SLIP-2d | 47.1 | 0.219 | – | 37.2 | 0.150 | – | 11.6 | 0.095 | – | 5.7 | 0.064 | – |
| ES-SLIP-d | 45.9 | 0.211 | – | 37.8 | 0.165 | – | 17.5 | 0.104 | – | 6.9 | 0.074 | – |
| SLIP | 70.3 | 0.264 | – | 57.5 | 0.185 | – | 55.0 | 0.122 | – | 41.2 | 0.088 | – |

**Table S13:** High versus low friction (Goldsmith et al.), ten bird–speed conditions present on both surfaces: medians, median paired difference (low *−* high) and Wilcoxon signed-rank *P* . Fitted values on the parameter grid (pivot step 0.15 *l*_0_).

|  | high | low | difference | *P* |
|---|---|---|---|---|
| speed (m s<sup>−1</sup>) | 1.71 | 1.70 | -0.03 | 0.01 |
| stance time (s) | 0.165 | 0.158 | +0.003 | 1.00 |
| peak vertical force (BW) | 1.76 | 1.75 | -0.00 | 0.56 |
| peak braking force (BW) | 0.266 | 0.282 | +0.021 | 0.08 |
| peak propulsive force (BW) | 0.208 | 0.229 | +0.028 | 0.02 |
| force-line intersection above COM (*l*<sub>0</sub>) | 0.57 | 0.63 | +0.05 | 0.38 |
| VPP2-SLIP-bl pivot height above COM (*l*<sub>0</sub>) | 1.52 | 1.00 | -0.15 | 0.06 |
| VPP2-SLIP-bl *ĉ*<sub>*bl*</sub> | 1.8 | 2.4 | +0.0 | 0.50 |
| RFd-VPP-SLIP-bl pivot height above COM (*l*<sub>0</sub>) | 0.85 | 0.77 | -0.15 | 0.08 |
| TD-SLIP-2d *ĉ* | 0.15 | 0.30 | +0.00 | 0.59 |
| TD-SLIP-2d *ĉ*<sub>*bl*</sub> | 1.2 | 0.6 | -0.6 | 0.22 |

